# ALOX12B overexpression in the skin drives inflammasome/Th17 signaling axis to promote inflammation in the mouse model and human patients

**DOI:** 10.1101/2025.06.09.658245

**Authors:** Suman Singh, Farhan Ahmed, Naireen Fatima, Cholleti Sai Nikhith, Harshavardhan Bhuktar, Rafiq Ahmad Khan, Sumbul Afroz, Srikanth Battu, Kakularam Kumar Reddy, Manaswani Jagadeb, Saima Naz, Neha Sharma, Aradhan Mariam Philips, Tina Priscilla, Vaibhav Vindal, M Jerald Mahesh Kumar, Srinivas Oruganti, Manojit Pal, Pallu Reddanna, Nooruddin Khan

## Abstract

Inflammation plays a pivotal role in the etiopathogenesis of chronic inflammatory skin diseases. However, the underlying mechanism remains unclear. Here, we employed Gene expression meta-analysis and clinical validation strategy to dissect the global architecture of immune dysregulation responsible for inflammatory conditions in the skin. Using such approaches, we identified a gene signature comprising of *ALOX12B*, which is significantly upregulated in psoriatic and atopic dermatitis patient skin samples, and correlates with increased levels of pathological IL-1β and Th17 responses. Surprisingly, *ALOX12B* is predominantly expressed in the skin. Furthermore, skin-specific overexpression of human *ALOX12B* in transgenic mice resulted in psoriasis-like inflammatory symptoms, including epidermal hyperplasia, immune cell infiltration, and elevated IL-1β/Th17 responses. ALOX12B is a non-heme iron-containing enzyme that catalyses the production of *12R*-HETE from polyunsaturated fatty acids such as arachidonic acid. Mechanistically, we found that ALOX12B/*12R*-HETE accumulation in the skin acts as an intrinsic danger signal that triggers enhanced IL-1β processing and secretion via ROS generation and NLRP3 inflammasome activation. Increased IL-1β levels in turn drive IL-17 producing T-cell polarization. We further designed a novel first-in-class, potent ALOX12B inhibitor, **6a**, which exhibited favorable topical pharmacokinetic and safety profiles. The topical application of **6a** reduced inflammation-associated pathologies in Tg-hALOX12B mice by suppressing *12R*-HETE-induced IL-1β production and ROS generation. These findings revealed a novel mechanism mediated by ALOX12B/*12R-*HETE overexpression that controls skin inflammation, thereby providing a promising therapeutic target for treating inflammatory skin diseases.

Graphical Abstract.
The proposed mechanism through which ALOX12B modulates the inflammatory pathway in psoriasiform-like inflammatory conditions in the skin.
The study provides insight into the role of ALOX12B in skin inflammation. Increased levels of ALOX12B expression in the skin result in 12R-HETE accumulation, which subsequently triggers the IL-1β/Th17 inflammatory cascade through ROS generation and inflammasome activation. IL-1β produced by the activated inflammasome leads to the polarization of naïve T-cells to activated IL-17 producing CD4^+^ T-cells, which are involved in the skin-inflammation related pathologies such as keratinocyte proliferation, and cellular infiltration. Hence, we propose ALOX12B as a novel potential therapeutic target for psoriasis-like inflammatory skin disorders. Additionally, we designed a novel first-in-class ALOX12B inhibitor, 6a; which shows a substantial alleviation in ALOX12B enzymatic activity, thereby reducing cellular ROS, which in turn results in reduced levels of IL-1β, as a result inhibiting IL-1β/Th17 inflammatory pathway and subsequent reduction in pathological inflammatory skin conditions. A portion of the image was created using Biorender.com

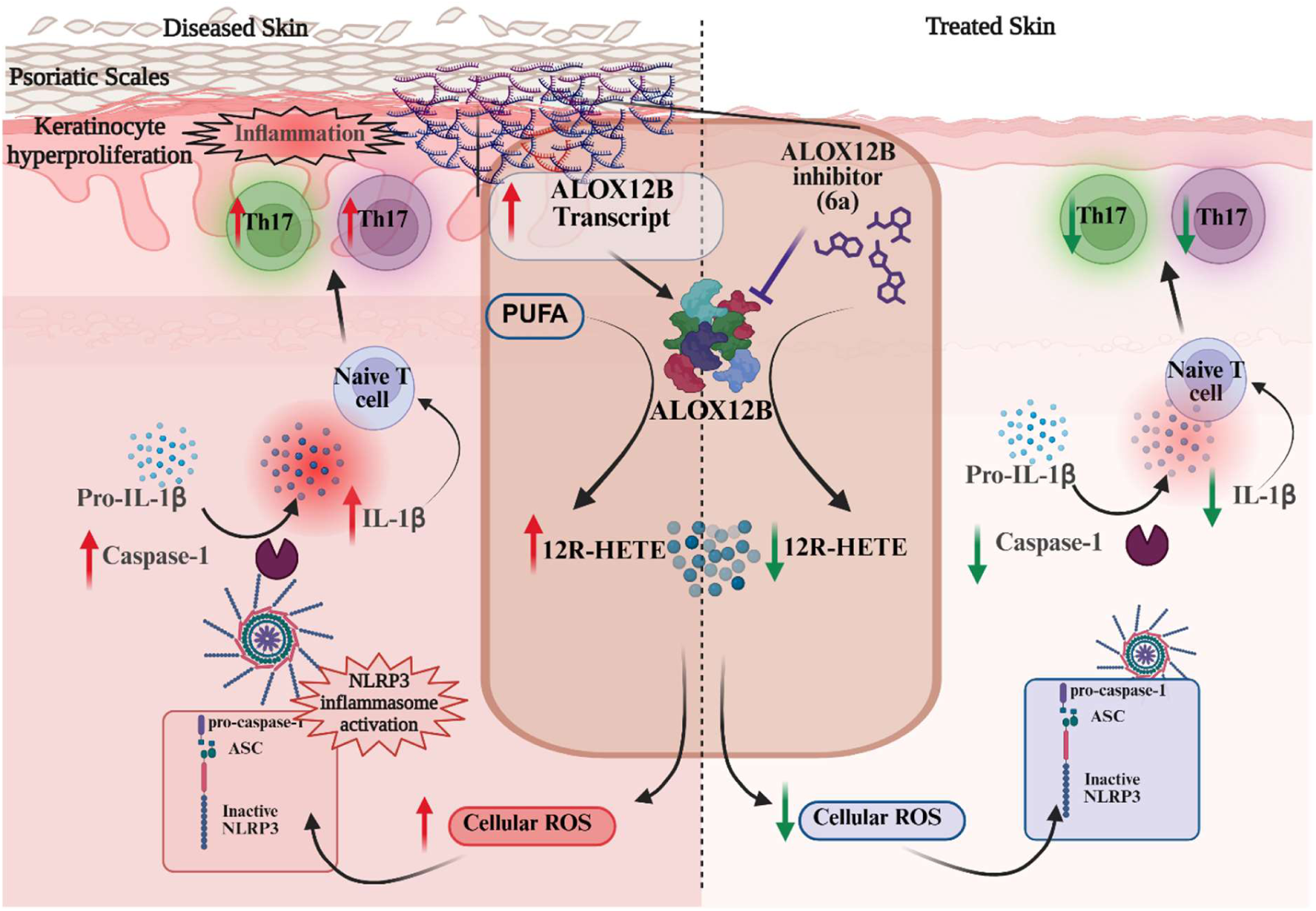

**Highlights:**

- ALOX12B is a key gene upregulated in the skin during psoriasis and atopic dermatitis.
- Transgenic mice overexpressing ALOX12B in the skin developed Psoriasis-like inflammatory symptoms.
- Mechanistically, ALOX12B, through *12R*-HETE, enhanced IL-1β production via ROS generation and NLRP3 inflammasome activation. Further elevated IL-1β production promotes Th17 cell polarization.
- Novel first-in-class ALOX12B inhibitor alleviates inflammatory symptoms in a transgenic mouse model.
- Study reveals a crucial ALOX12B/*12R-*HETE-inflammasome-IL-17 axis involvement in inflammatory skin diseases.

## Introduction

Psoriasis and Atopic dermatitis (AD) are prototypical chronic inflammatory skin diseases associated with the breakdown of skin homeostasis triggered by a multitude of factors such as genetics, environment, autoimmunity, and infections^1,2^. Collectively, they pose a serious challenge to public health, affecting 10 % of the adult population in the United States^3–5^. These debilitating and stigmatizing conditions are characterized by dysregulated keratinocyte proliferation, differentiation, dermal angiogenesis, and immune cell infiltration^6,7^ Although there is no permanent cure, advancements in biological therapies targeting specific dysregulated immune pathways, using antagonists for IL-17/IL-23 in psoriasis and JAK1/IL4Rα inhibitors in AD, offer a better treatment regimen. Despite remarkable advances, approximately 20-50 % of patients still do not achieve the desired outcomes with the current drugs^8–10^, underscoring the urgent need to deepen our understanding of the complex etiology of the disease to develop a more effective therapeutic regimen.

Recent advances in systems approaches have proven instrumental in identifying immune signatures associated with various immunological processes^11,12^. In line with these findings, we employed gene meta-analysis and clinical validation approaches to dissect the global architecture of the immune dysregulation involved in the pathogenesis of inflammatory skin diseases. A Meta-analysis was performed on publicly available gene expression datasets of Psoriatic patients, followed by clinical validation of the meta-gene profile in the skin biopsies of patients with psoriasis. Using such approaches, we identified a gene signature comprising of *ALOX12B,* which is significantly upregulated in all patient samples and correlated with increased levels of pathological IL-1β and Th17 responses, thereby suggesting ALOX12B could be a crucial target involved in the etiopathogenesis of psoriasis.

ALOX12B is a member of the non-heme iron-containing lipoxygenase (LOXs) family of enzymes which catalyzes the oxygenation of polyunsaturated fatty acids, such as arachidonic acid, which are crucial for various cellular functions^13^. In mammals, LOXs are expressed in multiple cell types, including epithelial, endothelial, and immune cells, and they play vital roles in cell function, differentiation, and immunity^13^. The human genome encodes six functional *LOX* genes (*ALOX15, ALOX15B, ALOX12, ALOX12B, ALOX5, and ALOXE3*), each encoding distinct LOX enzymes^14^.

Although the biological functions and clinical relevance of most LOXs have been extensively studied, ALOX12B, which converts arachidonic acid to 12*R*-HETE, especially in the skin, remains poorly understood. Unlike other *LOXs*, *ALOX12B* is exclusively expressed in skin epithelial cells^15^. Notably, rare mutations in the coding regions of *ALOX12B* cause autosomal recessive congenital ichthyosis (ARCI)^16–18^, and double knockout of this gene is lethal in mice due to impaired skin barrier function^19^, underscoring its critical role in skin development. Conversely, accumulation of 12*R*-HETE, a product of ALOX12B, has been associated with psoriasis and other inflammatory skin diseases^20–23^. However, the mechanism through which ALOX12B regulates skin inflammation remains unclear.

Given the importance of ALOX12B in the development and exacerbation of skin inflammation, this study was undertaken to elucidate the underlying mechanism through gene expression and immune correlation studies in skin biopsies of human patients and through the development of a genetic mouse model overexpressing human *ALOX12B* (hALOX12B). Surprisingly, humanized transgenic mice overexpressing *ALOX12B* (Tg-hALOX12B) displayed chronic inflammatory skin morphology with elevated levels of pro-inflammatory IL-1β cytokines and a pathological Th17 response akin to human inflammatory skin diseases. It is well documented that the production of IL-1β in immune cells depends on priming signals provided by the appropriate PAMPs and DAMPs^24–26^ and the activation signal required for the processing and release of mature IL-1β^27^. Processing of active IL-1β is primarily dependent on proteolytic cleavage of Pro-IL-1β protein by caspase-1, triggered by inflammasome activation in response to danger signals^28^. To our knowledge, this is the first study to elucidate a complex relationship between ALOX12B and IL-1β production. Mechanistically, we demonstrated that ALOX12B regulates IL-1β processing and production through its enzymatic product, *12R*-HETE. We further showed that *12R*-HETE enhanced IL-1β processing and secretion of mature IL-1β in LPS-primed macrophages through reactive oxygen species (ROS) generation and inflammasome activation. Enhanced *12R*-HETE mediated IL-1β levels drive pathological Th17 polarity, exacerbating skin inflammation. In addition, we identified a novel inhibitor targeting ALOX12B enzyme activity. Topical application of the ALOX12B inhibitor provided remarkable protection from inflammation-associated pathologies in a genetic mouse model, Tg-hALOX12B and cell-based ALOX12B overexpression systems by dampening IL-1β production through suppression of ROS generation and inflammasome activation. In conclusion, this study revealed a novel axis of skin inflammation involving ALOX12B and inflammasome activation. This discovery provides new insights into the pathogenesis of skin inflammation and identifies potential novel therapeutic targets for the treatment of inflammatory skin diseases.

### Methodology

The entire human sample collection process adhered to the ethical guidelines established by the Institutional Ethics Committees of the University of Hyderabad, India and the Apollo Institute of Medical Sciences and Research (UH/IEC/2019/210). Written informed consent was obtained from all participants, including the patients and healthy controls. All animal experiments were performed in accordance with the Institutional Animal Ethical Committee guidelines of the University of Hyderabad (UH/IAEC/NK/2023-1/43/R1), Anthem Biosciences Pvt. Ltd., and Palamur Biosciences Private Limited.

### Patient recruitment and Acquisition of skin biopsy specimens

Psoriatic skin biopsies were obtained from individuals diagnosed with Psoriasis according to the established clinical criteria. Participants who reported to Apollo Hospital, Hyderabad, India, were recruited for this study. Healthy control participants, defined as individuals with no reported history of dermatological or significant medical conditions, underwent skin biopsies.

### Reagents and plasmids

LPS (Sigma-Aldrich), Protease Inhibitor (Sigma-Aldrich), ATP disodium salt (Sigma-Aldrich), 12*R*-HETE, *12S*-HETE, *15S*-HETE (Cayman Chemical), Lipofectamine 3000 (Invitrogen), NAC (Sigma Aldrich), NaCl (Merck), Pam3CSK4 (InvivoGen), R848 (InvivoGen), poly(I:C) (InvivoGen), and MSU (InvivoGen). Mouse enzyme-linked immunosorbent assay (ELISA) kits for TNF-α (560478, BD Biosciences), IL-6 (Cat No.-550950, BD Biosciences), IL1-β (900-K47, PeproTech), and TMB Substrate Reagent Set (ER1200A, G Biosciences) were used. pCMV6-neo-ALOX12Bwas used for transfection studies, pBluescript SK (pBSK) was used to subclone the keratin 14 (K14) vector, pGE3Z.K14 (a gift from Elian Fusch from Rockefeller University, USA) was used for the development of hALOX12B overexpressing transgenic mice (Tg-ALOX12B), and pET28(b) was used for the cloning and expression of ALOX12B protein.

### Data Acquisition and preliminary processing

Publicly available RNA-sequencing (RNA-seq) and microarray gene expression datasets were systematically retrieved from the National Center for Biotechnology Information (NCBI) Gene Expression Omnibus (GEO)^29^. An initial collection of 10 datasets, specifically comprising human skin samples from psoriatic patients (including both lesional and non-lesional biopsies) and healthy control subjects, was identified. These datasets underwent a rigorous curation process to ensure the integrity of comparative analyses. This involved the distinct stratification of non-lesional psoriatic skin samples from healthy control samples before their final inclusion. This meticulous approach yielded 12 suitable datasets for the study (Table S1). Specific inclusion criteria further mandated that all samples originated from skin tissue and excluded potential confounding factors related to age, sex, race, and ethnicity.

To identify differentially expressed genes (DEGs) in each dataset, normalized gene expression values were compared between the lesional skin samples and healthy controls. Healthy controls included both non-lesional psoriatic skin and skin samples from healthy individuals. After stratifying samples into non-lesional skin from psoriatic patients and healthy controls, individual analysis of each dataset yielded 12 distinct sets of differentially expressed genes (DEGs) (Table S1). Each curated dataset was subjected to differential expression analysis using the R statistical environment^30^ and Bioconductor packages DESeq2^31^, limma^32^, and affy^33^. Log_2_fold changes were computed, and p-values were adjusted to control the false discovery rate (q-value) using the Benjamini-Hochberg procedure. Genes with q-values > 0.05 were filtered out. Gene annotation was performed using the annotate package.

### Meta-analysis

Each dataset was independently analyzed to identify differentially expressed genes (DEGs). A q-value threshold of 0.05 was used to define significant DEGs, while highly significant DEGs were designated as those with q-values lower than 0.000001 (Supplementary Data Fig. 3a). Following the creation of these two ranked lists for each dataset, we employed an intersectional strategy (as detailed in Fig. 1A) to establish the meta-gene expression profile (MGEP) and highly significant meta-gene expression profile (HSMGEP) associated with psoriasis.

**Fig 1.**
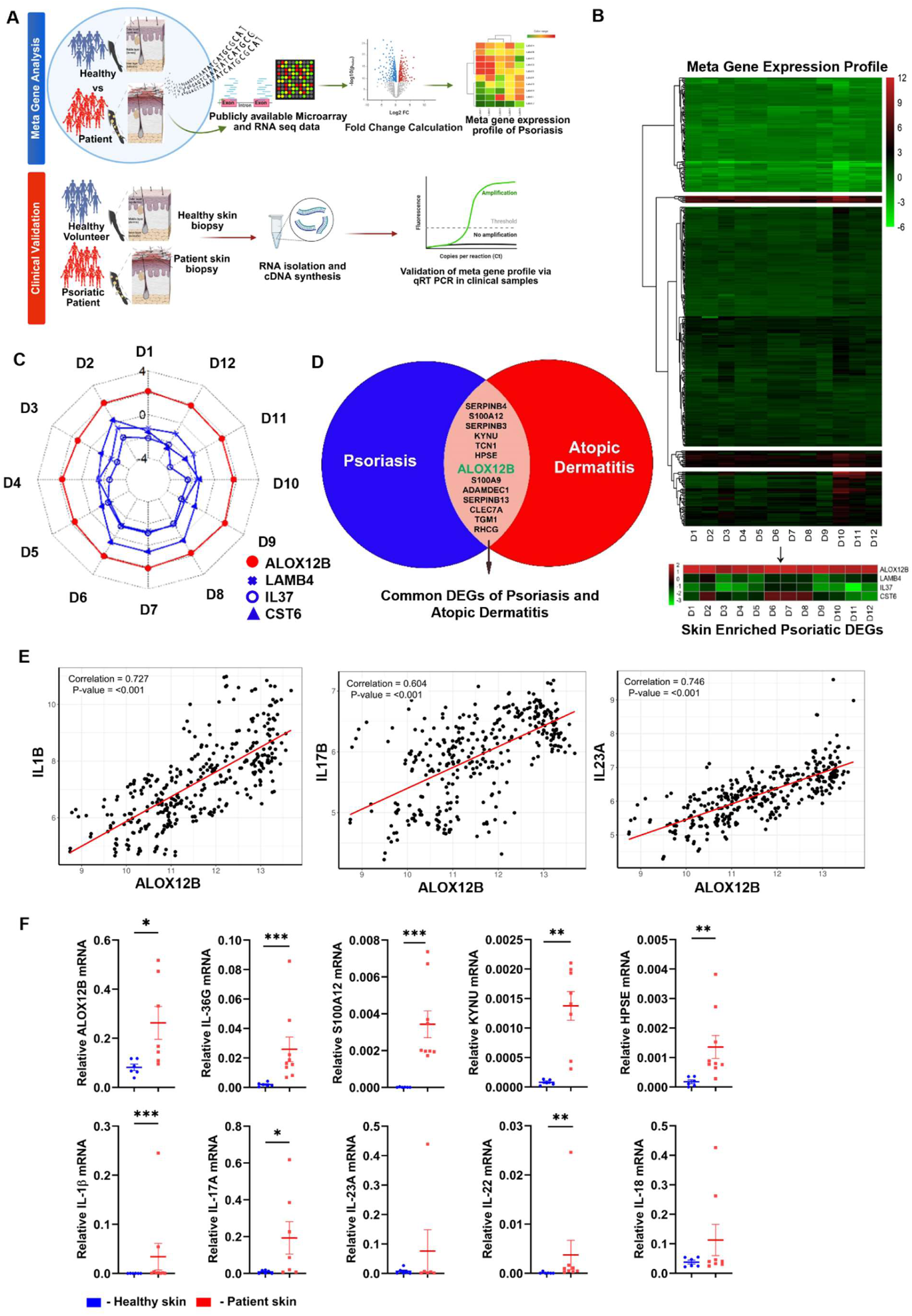
Multi-cohort meta-analysis and clinical validation approach in human patients identified *ALOX12B* as a key immune signature associated with inflammatory skin diseases. **A)** Schematic representation of the study design, outlining the integration of systems-level immunological analysis with subsequent clinical validation on human skin samples. The upper panel depicts the meta-analysis workflow employed to identify differentially expressed genes (DEGs) in psoriasis, utilizing publicly available datasets from the Gene Expression Omnibus (GEO). The lower panel illustrates the experimental methodology for quantitative real-time polymerase chain reaction (qRT-PCR) analysis of gene expression profiles, performed on skin biopsy samples procured from individuals diagnosed with psoriasis and healthy control subjects. **B)** The upper panel displays a heatmap visualization of the MGEP, encompassing 435 statistically significant DEGs identified through meta-analysis of 12 curated psoriasis datasets derived from the NCBI Gene Expression Omnibus (GEO). Hierarchical clustering, utilizing Euclidean distance for dissimilarity measurement and complete linkage, was applied to the MGEP, resulting in the segregation of genes into five distinct expression pattern-based clusters. The lower panel presents a heatmap of the four skin-enriched psoriatic DEGs (SEPDs) obtained by intersecting the MGEP with a set of skin-enriched genes. The color intensity scale indicates the relative expression levels of genes. **C)** Spider plot illustrating the log2 fold change of four skin-enriched psoriatic DEGs (SEPDs) across 12 independent datasets. The red lines represent genes upregulated in psoriasis, while the blue lines represent downregulated genes. **D)** Venn diagram depicting the overlap in dysregulated genes identified in psoriasis and atopic dermatitis. The intersecting area highlights a subset of commonly altered genes, specifically on ALOX12B, suggesting its potential significance in the shared molecular pathology of these inflammatory skin conditions. **E)** Pairwise scatter plot and correlation analysis of ALOX12B and selected pro-inflammatory markers IL1B, IL17B, and IL23A. Each scatter plot displays the normalized expression values of the indicated gene pair, along with the corresponding Pearson correlation coefficient (r) and p-value. The data reveal a positive correlation between ALOX12B expression and the expression levels of these key inflammatory cytokines in the analyzed datasets. **F)** Quantitative real-time PCR (qRT-PCR) analysis was performed on skin biopsy samples obtained from individuals diagnosed with psoriasis and healthy control subjects to validate the differential expression of key genes identified through meta-analysis. Statistical significances were determined by two-tailed unpaired t-test; \**P* ≤ 0.05; \*\**P* ≤ 0.01; \*\*\**P* < 0.001. Data are shown as means ± SEM.

### Gene Ontology (GO) and pathway enrichment

Functional enrichment analysis of the identified MGEP was performed to elucidate its biological relevance. The Enrichr web-based tool^34,35^, specifically designed for pathway and Gene Ontology (GO) enrichment analysis, was utilized to predict enriched GO terms in biological processes, molecular function, and cellular component categories.

### Quantitative real-time PCR (qRT-PCR) of human skin samples

Total RNA was isolated from skin biopsy samples preserved in RNA *later* solution (AM7020, Invitrogen) using an RNA extraction kit (74134, Qiagen) following the manufacturer’s protocol. The purified RNA was reverse-transcribed into complementary DNA (cDNA) using a cDNA synthesis kit (AB-1453/A, Thermo Scientific) according to the manufacturer’s instructions. Approximately 40 ng of cDNA was used as a template in the qRT-PCR reaction mixture. cDNA was subsequently amplified using SYBR Green master mix (RR820A, TAKARA) and gene-specific primers (Table S13). The Thermal cycling conditions included an initial denaturation step at 95 °C for 5 min, followed by 40 cycles of amplification consisting of denaturation at 95 °C for 30 s, annealing at 60 °C for 30 s, and extension at 72 °C for 40 s. Relative mRNA expression levels for each sample were determined by normalization to the housekeeping gene GAPDH.

### Generation of human *ALOX12B* overexpressing transgenic mice

The pCMV6-Neo-hALOX12B plasmid was procured from OriGene (USA), containing complete cDNA along with the 5’ and 3’ UTR sequences of human *ALOX12B*. For skin-specific overexpression of hALOX12B, *ALOX12B* cDNA from pCMV6-Neo-hALOX12B was released by NotI digestion and subcloned into the pBKS plasmid by blunt end ligation at the SmaI site to generate Xba1 sites on either side of *hALOX12B*. From pBKS-*hALOX12B*, whole cDNA was released by XbaI digestion and ligated into pGE3Z.K14 (a gift from Elian Fusch from Rockefeller University, USA). For the generation of *hALOX12B* overexpressing transgenic mice specifically in the skin, the pGE3Z.K14-hALOX12B vector was double digested with SmaI/HindIII to release the whole cassette (5.7 kb) containing the K14 promoter, beta-globin intron, human *ALOX12B* cDNA, and polyA tail. This fragment was then purified and microinjected into mouse embryos. Gene integration and germline transmission of the transgene were confirmed by PCR using gene-specific primers and Southern blotting, respectively. Skin biopsies from litter mate controls and transgenic (Tg-hALOX12B) mice were used to determine the transcript and protein levels of *hALOX12B* in Tg-hALOX12B. Southern blot analysis was performed by digestion of mouse genomic DNA with XbaI or BamHI. The fractionated DNA was run on an agarose gel and transferred onto a hybond-N^+^ membrane using a VacuGene XL Vacuum Blotting system according to the manufacturer’s instruction or as described by Southern^36^. DNA was radiolabeled with the probe and scanned using a phosphorimager (Fujifilm).

### Genotyping of Tg-hALOX12B mice

PCR analysis was performed for the selection of positive clones with Tg-hALOX12B. Briefly, tail biopsies from mice were digested overnight in tail lysis buffer (1M Tris-HCl, 5M NaCl, 0.5M EDTA, 10 % SDS, and 20 mg/ml proteinase K) at 57 °C. The lysis supernatant was extracted after centrifugation at 12000 rpm, and an equal amount of Tris-saturated phenol was added. The supernatant was again extracted after centrifugation at 12000 rpm, and an equal volume of phenol:chloroform:isoamyl alcohol (25:4:1) was added. Similarly, equal volumes of chloroform and isopropanol were sequentially added to the supernatant. Finally, genomic DNA was precipitated in isopropanol by centrifugation, pellet was washed with 70 % ethanol and resuspended in Tris-EDTA buffer pH 8 (as described previously with modifications)^37^. Genomic DNA (100 ng of genomic DNA was used for each PCR amplification using hALOX12B specific primer for genotyping (Table S13) with the following parameters: one cycle of initial denaturation at 94 °C for 10 min. This was followed by 35 cycles of denaturation for 30 s at 94 °C, primer annealing for 30 s at 57 °C, and extension for 2 min 30 s at 72 °C. The amplified product was observed by gel electrophoresis for the positive clones of Tg-hALOX12B mice.

### qRT-PCR

For transcript-level expression, RNA was isolated using TRI reagent (T9424, Sigma-Aldrich). The purified RNA was reverse-transcribed to cDNA using a cDNA synthesis kit (6110A, TAKARA) according to the manufacturer’s protocol, and 50 ng of cDNA was used for each qRT-PCR reaction. qRT-PCR was performed on an Applied Biosystems thermal cycler (Thermo Fisher Scientific). The reverse-transcribed cDNA was amplified with SYBR green (RR820A, TAKARA) using primers for specific genes (Table S13) on the following thermal cycler parameters: one cycle of initial denaturation at 95 °C followed by 40 cycles of denaturation for 15 s at 95 °C, primer annealing for 30 s at 60 °C, and extension for 30 s at 72°C.

### Skin Histopathology

Tg-hALOX12B mice with phenotypically visible lesions of hair loss patches and wild-type control mice aged 9 to 12 months were used. Tissues fixed in 4 % paraformaldehyde, embedded in paraffin, and sectioned using a microtome. The Paraffin sections were deparaffinized with xylene and then rehydrated by sequential washing with ethanol (100 %, 90 %, 70 %and 50 %) and PBS for 5 minutes each^38^. Tissue sections were subjected to H&E (Haematoxylin and Eosin) staining or Masson’s trichrome staining.

### Haematoxylin and Eosin (H&E) staining

For H&E staining, deparaffinized sections were stained with hematoxylin, washed, and stained with eosin, followed by sequential dehydration with a graded series of ethanol and xylene^39^. The sections were mounted using a mounting agent and visualized under a light microscope.

### Masson’s trichrome staining

For Masson’s trichrome staining, the deparaffinized sections were sequentially stained with Weigert’s iron hematoxylin, Biebrich scarlet-acid fuchsin solution, phosphomolybdic-phosphotungstic acid solution, and aniline blue solution. Thereafter, rinsed with distilled water and 1 % acetic acid, followed by sequential dehydration with graded series of ethanol, then with xylene^40^. The sections were mounted using a mounting agent and visualized under a light microscope.

### Immunohistochemistry

For immunohistochemistry, deparaffinized sections were subjected to antigen retrieval with 10 mM sodium citrate (pH 6) at the boiling temperature for 5 min, followed by cooling. To minimize the non-specific binding of antibodies, sections were blocked with 10 % goat serum or BSA for 1 h at room temperature, followed by incubation with primary antibody overnight at 4 °C. After washing, the sections were incubated with Alexa Flour-conjugated or HRP-conjugated goat anti-mouse (Invitrogen, USA) secondary antibody for 1 h at room temperature. The primary antibodies used included E-selectin (CD62E, BioLegend), VCAM 1 (CD 106 BioLegend), PECAM (CD38, BioLegend), CD4, CD8, Col 1, and MPO. Sections stained with horseradish peroxidase (HRP)-conjugated antibodies were developed using a chromogen. All sections were washed, and coverslips were mounted with an anti-fade medium (Vectashield, Vector Laboratories) and examined under a trinocular fluorescent microscope (Leica Microsystems). Images were captured using Leica software, overlaid, and processed using imageJ2 (fiji), an open-source software from the NIH.

### Flow cytometry analysis of mouse spleen, and PBMCs

Spleen was mechanically dissociated, lysed with RBC lysis buffer, and filtered sequentially with 70- and 40-μm nylon mesh filters. PBMCs were isolated by sucrose gradient density separation using Histopaque (Sigma-Aldrich). For T cell intracellular cytokine staining, cells were stimulated with 50 ng/mL phorbol myristate acetate (p1585, Sigma-Aldrich) and 1 µg/mL ionomycin (I0634, Sigma-Aldrich) in the presence of Golgi stop (BD Bioscience) for 8 h before surface staining with antibodies. Isolated cells were stained in FACS buffer (PBS, 3 % FBS, and 2 mM EDTA) with the following antibodies; PerCP labeled anti-mouse CD4, FITC-labeled for 1 h. For the intracellular staining, cells were washed three times, fixed with 4 % paraformaldehyde, permeabilized with 1x perm wash buffer (Biolegend), and stained with phycoerythrin-conjugated anti-mouse IL-17 antibody diluted in perm wash buffer for 1 h at room temperature. Cells were washed again and resuspended in FACS buffer, and data were acquired on a BD Accuri C6 or BD LSR Fortessa cytometer (BD Biosciences) and analyzed using FlowJo software 10.10.0.

### Cell Culture and Treatment

J774A.1, procured from the National Center for Cell Science (NCCS), Pune, India, was maintained in DMEM (Gibco, Invitrogen), supplemented with 10 % FBS (v/v) (Gibco, Invitrogen), penicillin (100 units/ml), streptomycin (100ug/ml) (Gibco, Invitrogen), 2mM L-Glutamine (Gibco), 1mM sodium pyruvate (Gibco, China) at 37 ℃ with 5 % CO_2_ in a humidified incubator. Macrophages were primed with LPS (500ng/ml), Pam3CSK4 (1 μg/ml), R848 (10 μg/ml), poly(I:C) (10 μg/ml), or MSU (100 ug/ml) for 5 h and then stimulated with 1µM µM *12*R-HETE, *12S*-HETE, *15S*-HETE, or MSU (500 μg/ml) for 1 h or ATP (5 mM) for 30 min. In some experiments, LPS-primed macrophages were treated with NAC (10 mM), KCl (130 mM), or NaCl (130 mM) for 2 h and then with *12*R-HETE for 1 h. For the *in vitro* screening of ALOX12B inhibitor, **6a** macrophages were transfected with pCMV6-hALOX12B. After 18 h of incubation, the cells were treated with LPS (500 ng/ml) and **6a** (10 μM) for 6 h.

### Cell Transfection

*In vitro* transfection was performed using Lipofectamine 3000 according to the manufacturer’s instructions. Briefly, J774a.1 cells at 70 % confluency in complete media were transfected with pCMV6-neo-hALOX12B or empty vector control. The cells were then incubated for 18 h for further analysis and treatment.

### Generation of mouse bone marrow-derived macrophages

Bone marrow from the femur and tibia of C57BL/6 mice was isolated, and the cells were flushed in RPMI 1640 medium. Cells were cultured at a density of 5 × 10^6^ or 7 × 10^6^ in 100 mm Petri dishes supplemented with DMEM with 10 % FBS (v/v), penicillin (100 units/ml), and streptomycin (100 µg/ml) in the presence of 20 ng/ml MCSF (Peprotech). Fresh medium containing MCSF was added on Day 4. After 6 days of incubation, the cells were harvested for further experiments^41^.

### Immunoblotting

For protein expression analysis, skin tissue was crushed in liquid nitrogen and lysed in RIPA buffer (Thermo Scientific) supplemented with protease inhibitor cocktail tablets (Sigma-Aldrich) according to the manufacturer’s protocol. Cells were harvested, washed with PBS, and lysed in lysis buffer. The lysate was incubated on ice for 40 min with occasional vortexing, and a clear lysis supernatant was obtained after centrifugation at 12000 rpm for 20 min at 4 °C. Protein concentration was estimated using Bicinchoninic Protein Assay (BCA) kit (G Biosciences) as according to the manufacturer’s instructions, and equal concentrations of protein were loaded per well and separated on a 12 % SDS-PAGE gel. Proteins were transferred onto a nitrocellulose membrane by electroblotting, and the membrane was blocked with 5 % skim milk in PBS for 1h at room temperature. Antibodies used for immunoblotting were Anti-ALOX12B (Sigma-Aldrich ATLAS Antibody), IL-1β (Santa-Cruz or Gentech), Caspase-1 p10 (Santa Cruz or Adipogen), and β-Actin (Cell Signaling Technology or Santa Cruz) as loading control. Primary antibodies were incubated at 4 °C overnight, then 3 times washed with PBST and incubated with appropriate HRP-tagged secondary antibody for 1 h at room temperature and again washed 3 times with PBST to remove non-specifically bound antibodies. Bands were then observed with a chemiluminescence substrate (G Biosciences), and chemiluminescence was observed in the ChemiDoc Touch Imaging System (Bio-Rad).

### Measurement of Cleaved IL-1𝛽

Macrophages with or without treatment, cultured in Opti-MEM, were stimulated with LPS (500 ng/ml) for 6 h. Post-treatment, the cell supernatant was harvested, and protein precipitation was performed following the methanol-chloroform method as described earlier^41^. Briefly, 500 µl of supernatant was mixed with equal amounts of methanol followed by the addition of 100 µl of chloroform, vortexed vigorously, and centrifuged at 12000 rpm for 5 min. The aqueous layer was removed, and 500 µl of methanol was added, vortexed again, and centrifuged at 14000 rpm for 2 min. The supernatant was discarded, and the pellet was dried at 50 ℃ for 5 min and dissolved in 30 µl of 2x Laemmli sample buffer (Bio-Rad). The sample in sample buffer was heated at 95℃ for 5 min, sample was loaded and separated on a 12 % SDS-PAGE gel. Protein was transferred onto the nitrocellulose membrane by electroblotting and cleaved IL-1𝛽 bands were measured by immunoblotting as described in the previous section.

### Cytokine ELISA

Cytokines in the cell culture supernatants of the different treatment groups were measured using sandwich ELISA. IL-1β, IL-6, and TNF were measured as per the manufacturer’s instructions. Briefly, the polystyrene 96 well plate was coated with anti-cytokine antibody overnight at room temperature for IL-1β and 4 °C for IL-6, TNF-α. Plates were washed with PBST and blocked with 1 % BSA at room temperature for 1 h, followed by incubation with the test sample. Plates were then washed again and incubated with detection anti-cytokine antibody conjugated with biotin followed by incubation with horse-reddish peroxidase conjugated with streptavidin. TMB substrate (G Biosciences) was used for the reaction. The reaction was stopped by 1 N H_2_SO_4_ and absorbance was measured at 450 nm and wavelength correction 562 nm^42^.

### ROS detection

For ROS estimation, macrophages were treated with LPS (500 ng/ml) for 5 h, HETE (1µM/ml) for an additional 1 h. Cells were extracted, washed with HBSS, and incubated with CM-H_2_-DCF-DA (I36007, Life Technologies) for 30 min at 37°C. The cells were then washed again with HBSS^41^. ROS levels were estimated using flow cytometry. Data were acquired on a BD Accuri C6 cytometer (BD Biosciences) and analyzed using FlowJo software 10.10.0.

### Immunofluorescence Microscopy

Macrophages were seeded and incubated overnight on a coverslip in 24 well plate, followed by appropriate treatment with LPS (500 ng/ml) for 5 h, and then 12*R*-HETE (1 µM/ml) for an additional 1 h. Cells were fixed with 4 % paraformaldehyde for 15 min at room temperature, washed three times with 1x PBS for 5 min each wash, and permeabilized with 0.2 % Triton X-100 for 20 min, washed as mentioned previously. Later, cells were blocked with 5 % BSA for 1 h at room temperature and washed again. Cells were incubated with primary antibody for 2 h at room temperature or overnight at 4°C, followed by secondary antibody incubation tagged with Alexa Fluor 555 (Invitrogen). The cells were mounted with an anti-fade medium (Vectashield, Vector Laboratories). Images were captured under the Leica confocal microscope and analyzed using Image J2 (Fiji), an open-source software from the NIH.

### *In vitro* screening of ALOX12B inhibitor

*ALOX12B* was fused with an N-terminal His-tag cloned into the pET28b vector and expressed in Rossetta cells (BL21). The cells were allowed to grow at 37 °C, IPTG (1 mM) was added to the culture when it reached an O.D of 0.5-0.6. The culture was shifted to 18 °C after IPTG induction. After 10-12 hrs., the cells were harvested by centrifugation at 6000 rpm at 4 °C for 10 min. The cells were washed with PBS (Buffer) and lysed in PBS containing lysozyme (conc.) and protease inhibitor cocktail (Rosch). The lysed cells were centrifuged at 13000 rpm for 30 min at 4 °C to obtain a clear supernatant containing the soluble recombinant protein. Purification of the recombinant protein was performed using Ni-NTA affinity chromatography according to the manufacturer’s protocol. Purified hALOX12B protein was added to a total volume of 0.5 ml of PBS containing arachidonic acid methyl ester as a substrate at a final concentration of 0.1 mM. The mixture was vortexed and incubated for 15 min at 37 °C, and the hydroperoxy fatty acids formed were reduced to more stable hydroxy derivatives by the addition of sodium borohydride saturated solution. After 5 min, the reaction was acidified to pH 3 with 50 μl of acetic acid, and the proteins were precipitated with the addition of 0.5 ml of methanol. The total mixture was centrifuged at 14000 rpm for 15 min to remove proteins, and the clear supernatant was analyzed by reverse-phase HPLC (RP-HPLC), using C-18 column. High-pressure liquid chromatography was performed on a Shimadzu system equipped with a Hewlett-Packard diode array detector 1040A and HP-Chemstation program. The solvent system used was methanol: water: acetic acid in a ratio of 80:20:0.1 with a flow rate of 1 ml/min^43^.

### Metabolic stability of 6a in mouse liver microsomes

An *in vitro* metabolic stability study for **6a** at 5 μM test concentration in mouse liver microsomes (0.25 mg/mL) was performed to determine the half-life (t1/2, min). *In vitro* intrinsic clearance (CLint, in vitro μL/min/mg of protein) was performed using mouse liver microsomes. The detailed procedure is provided in the supplementary methods.

### Intravenous pharmacokinetic (IV) profile of 6a

To evaluate the plasma pharmacokinetic profile of the test compound **6a**, male BALB/c mice were administered an intravenous dose at 10 mg/kg body weight. Plasma concentrations of **6a** were quantified by LC-MS/MS, and pharmacokinetic parameters were calculated as described in the Supplementary Methods.

### Single Dose Dermal Pharmacokinetic Study of 6a in Mice

10 mg/ml of **6a** dissolved in DMSO was applied to hair cleaned mice for 24 h. Blood and skin were collected for further analysis, as described in the supplementary methods.

### Acute dermal toxicity study of 6a in Wistar rats

The “Acute Dermal Toxicity” study was conducted as per Organization for Economic Co-operation and Development Guideline for Testing of Chemicals “Acute Dermal Toxicity: Fixed Dose Procedure” (Section-4, No. 402; adopted 9th October 2017), as described in the supplementary methods.

### *In vitro* hERG channel (ikr) assay

The inhibitory effect of **6a** on the rapid component of the delayed rectifier potassium current (IKr) was investigated using Human Embryonic Kidney cells stably transfected with hERG, employing the whole-cell patch-clamp technique, as described in the supplementary methods.

### Topical application of the ALOX12B inhibitor 6a in transgenic mouse model

The ALOX12B inhibitor 6a was topically applied at a dose of 50 mg/kg body weight to the visible hair loss patches of Tg-hALOX12B mice. Briefly, the compound was dissolved in DMSO, homogenized with a petroleum jelly base, and applied daily for 10 days to the affected skin patches of Tg-hALOX12B mice.

### Statistical analysis description

Statistical analyses were performed using GraphPad Prism 10 (GraphPad Software Inc.). For the comparison of two groups, Student’s t-test was used; when comparing three or more groups, Tukey’s test (one-way ANOVA) was used. All data are shown as mean values, and error bars represent the standard error of the mean (SEM) from three or more independent assays and are plotted in the graph. *P* values are indicated as \**P* < 0.05, \*\**P* < 0.01, \*\*\**P* < 0.001, *P* > 0.05, not significant (ns).

### Chemistry

#### General methods

Reactions were monitored by thin-layer chromatography (TLC) on silica gel plates (60 F254) and visualized with ultraviolet light or iodine spray. ^1^H and ^13^C NMR spectra were recorded in CDCl3 or DMSO-d6 using 400 and 100 MHz spectrometers (VARIAN 400 MR), respectively. The proton chemical shifts (δ) were relative to tetramethylsilane (TMS, δ = 0.00) as an internal standard and are expressed in ppm. Spin multiplicities are given as s (singlet), d (doublet), t (triplet), m (multiplet), and the coupling constants (*J*) are given in hertz. The melting points were determined using a melting point apparatus (Buchi melting point B-540) and were uncorrected. Mass spectrometry (MS) spectra were obtained using a mass spectrometer (AGILENT 6430 triple quadrupole LC-MS). Chromatographic purity was determined by HPLC (Agilent 1200 series Chem Station software) using the area normalization method and the conditions specified in each case: column, mobile phase (range used), flow rate, detection wavelength, etc. A schematic representation of the procedure for chemical synthesis is described in the supplementary methods.

## Results

### Multi-cohort meta-analysis and clinical validation approach in human patients identified *ALOX12B* as a key immune signature associated with inflammatory skin diseases

The complex etiology and heterogeneous clinical presentation of inflammatory skin diseases, such as psoriasis, necessitate a comprehensive systems-level investigation of global gene expression profiles and their immunological underpinnings to elucidate the core molecular drivers responsible for initiating and perpetuating chronic cutaneous inflammation. To address this, we systematically examined publicly available microarray and RNA-seq gene expression datasets pertaining to psoriasis.

Leveraging gene expression data derived from skin biopsies of 312 individuals with psoriasis and 389 healthy control subjects across 12 meticulously curated and independent datasets (Table S1), we performed a meta-analysis to identify consistently differentially expressed genes (DEGs). For each dataset, we calculated the log_2_ fold change and the corresponding q-value for every gene by comparing psoriatic skin to healthy or control skin. To identify the genes exhibiting robust and reproducible alterations across these studies, we determined the overlap of genes identified as significantly differentially expressed (at a q-value threshold of 0.05) within the 12 datasets. This rigorous selection process yielded a core set of 435 genes, which we designated as the “meta-gene expression profile” (MGEP). The identified MGEP represents a consensus set of genes that exhibit significant transcriptional dysregulation in psoriasis across diverse patient cohorts (Fig. 1B).

To prioritize genes within MGEP that are particularly relevant to skin biology and pathology, we filtered the 435 MGEP genes for those known to exhibit enriched expression in skin tissue. This was achieved by integrating our MGEP findings with skin tissue-specific gene expression data from the Human Protein Atlas database^44^. This intersectional analysis revealed four genes that were differentially expressed in the psoriasis meta-analysis and demonstrated preferential expression in skin tissue (Fig. 1B; lower panel). These four genes constitute a refined set, termed “skin-enriched psoriatic DEGs” (SEPDs). The four SEPDs identified were *ALOX12B*, *LAMB4*, *IL37*, and *CST6*. Notably, *ALOX12B* was consistently upregulated in psoriatic skin across the datasets included in the meta-analysis, whereas *LAMB4*, *IL37*, and *CST6* were consistently downregulated in most of the datasets. The log_2_ fold-change values for these four key SEPDs, demonstrating their expression patterns across the 12 independent psoriasis datasets, are shown in Figure 1C, highlighting the consistency of their dysregulation.

To assess the broader relevance of the identified MGEP signature in other common inflammatory skin conditions, we examined the expression status of the MGEP genes in atopic dermatitis (AD). We compared the MGEP genes against gene expression data from AD patients in the Expression Atlas database^45,46^. Comparative analysis revealed a significant overlap in the dysregulated genes between psoriasis and atopic dermatitis (AD), suggesting shared molecular pathways in skin inflammation. Specifically, a subset of genes differentially expressed in psoriasis within MGEP also exhibited differential expression in AD. Figure 1D illustrates some of these overlapping genes. Notably, *ALOX12B* expression was significantly elevated in the AD samples, mirroring our observations in psoriasis, indicating its potential importance in both conditions. Given that *ALOX12B* is known to be involved in skin barrier function and its enzymatic products, 12R-HETE has been reported to accumulate in psoriatic plaques^19,21,47^ its consistent and significant upregulation across multiple independent psoriasis datasets coupled with its significant upregulation in AD^48^ strongly suggests a pivotal role for *ALOX12B* in the broader context of chronic skin inflammation.

Further analysis of the log_2_ fold changes across the individual psoriasis datasets confirmed a consistent upregulation of *ALOX12B*, alongside key pro-inflammatory cytokines, including *IL1B*, *IL17A*, and *IL23A* (Supplementary Data Fig. 1). Moreover, pairwise correlation analysis revealed a significant positive correlation between the expression levels of *ALOX12B* and the central inflammatory mediators (*IL1B*, *IL17B*, and *IL23*) within the datasets (Fig. 1E). This correlation suggests a potential role for *ALOX12B* in regulating the inflammatory cytokine milieu, characteristic of psoriatic skin.

Gene Ontology (GO) enrichment analysis performed on the 435 genes comprising MGEP revealed significant enrichment directly associated with skin-related diseases and inflammatory responses. The enriched pathways, biological processes, molecular functions, and human phenotypes identified by GO analysis were predominantly linked to immune system processes, immune responses, immune cell function, infections, and skin-related infections or disorders (Supplementary Data Fig. 2a, 2b, 2c, and 2d). These findings underscore the crucial role of MGEP genes in orchestrating and modulating immune responses in cutaneous inflammation. Collectively, these system-level analyses robustly implicate MGEP genes in the development and progression of inflammatory skin conditions.

To further refine our findings and identify the most robust and consistently dysregulated genes, we applied a more stringent significance threshold (q-value < 0.000001), selecting DEGs that were highly significant across all the datasets. This analysis resulted in a refined set of 42 highly significant and reproducible DEGs, termed the “highly significant meta-gene expression profile” (HSMGEP) (Supplementary Data Fig. 3a and 3b). HSMGEP represents a particularly robust and consistently altered gene signature observed across all 12 datasets in our meta-analysis. Notably, *ALOX12B* is the part of the HSMGEP, further highlighting its significance in skin inflammation.

To investigate the tissue specificity of these highly significant genes, we analyzed their normalized expression levels (measured in Transcripts Per Million, TPMs) across a panel of various human organs using the Human Protein Atlas database. This analysis revealed a striking specificity of *ALOX12B* expression in the skin, with significantly higher transcript levels observed in the skin tissue than in other organs (Supplementary Data Fig. 4a). Interestingly, although *ALOX12B* exhibited this preferential expression pattern in the skin, other members of the arachidonic acid lipoxygenase family demonstrated broader expression patterns across different tissues (Supplementary Data Fig. 4b), highlighting the unique enrichment of *ALOX12B* in the skin context.

Based on the robust and significant findings from the systems approach, we further validated the expression of *ALOX12B* and other HSMGEP genes in skin biopsies of human subjects. Skin biopsies were obtained from 12 patients with psoriasis and 6 healthy volunteers using the standard skin biopsy punch method. A qualified dermatologist at Apollo Hospital, Hyderabad, India, performed all sample collections. The biopsies were quartered and frozen in RNA later for further analysis. The expression of prominent HSMGEP genes was validated by qRT-PCR. Notably, the validation confirmed a significant upregulation of *ALOX12B*, alongside key inflammatory mediators such as *IL1B*, *IL36G*, *IL22,* as well as genes associated with skin tissue reorganization including *KYNU, S100A12, and HPSE* (Fig. 1F). These genes are known to play critical roles in the pathogenesis of inflammatory skin diseases^49–53^. However, the immunomodulatory role of *ALOX12B* in skin remains poorly characterized. Notably, the prominent upregulation of *ALOX12B* in skin, along with its strong correlation with major inflammatory cytokines such as IL-1B, IL-22, IL23, suggests a crucial role of *ALOX12B* in modulating the etiopathogenesis of inflammatory skin disease.

### Overexpression of human *ALOX12B* in mice demonstrated a skin abnormality mimicking psoriasis-like skin inflammatory condition

Our meta-analysis and clinical validation approaches have revealed the importance of *ALOX12B* in inflammatory skin diseases such as Psoriasis and AD. Notably, *ALOX12B* predominant expression over other lipoxygenases in the skin further adds to its crucial role in skin pathobiology. Therefore, to elucidate the complete mechanism(s) through which *ALOX12B* overexpression leverages inflammatory mediators and tailors’ inflammation-associated pathologies in the skin, we developed a humanized transgenic mouse model expressing the human *ALOX12B* (Tg-hALOX12B) gene and studied its immunophenotypic characteristics. To develop Tg-hALOX12B mice, a construct containing the human *ALOX12B* cDNA under mice keratin-14 (K-14) promoter was injected into the mouse embryo. The mouse progenies were screened for the transgene, followed by validation of its integration and expression, as described in the supplementary data (Supplementary Data Fig. 5 and Table S2). Notably, the developed transgenic mice showed age-related phenotypic changes in skin morphology, including the appearance of patches and hair loss, compared to their wild-type littermates (Fig. 2A). Histopathological analysis of the skin revealed anatomical abnormalities in the skin of Tg-hALOX12B mice, such as thickening of the epidermis, epidermal hyperplasia, and infiltration of widespread infiltrates in the dermis region, which are the hallmarks of psoriasis-like skin inflammatory conditions (Fig. 2B upper panel). Interestingly, Masson’s trichrome staining of the skin sections demonstrated hyperkeratosis (dark red stain) and dermal fibrosis (blue stain) in the skin of Tg-hALOX12B mice compared to littermate controls (Fig. 2B lower panel). Moreover, measurement of histopathological parameters, including epidermal thickness, infiltrating cells, and cumulative histopathological score as seen by visual inspection in H&E-stained sections and Masson’s trichrome staining, demonstrated exacerbated inflammation-associated skin pathology in Tg-hALOX12B mice (Fig. 2C). We observed significant expression of the human *ALOX12B* gene in the skin of Tg-hALOX12B mice through qRT-PCR (Fig. 2D) and confirmed its enhanced protein level expression in the skin via western blotting (Supplementary Data Fig. 6), which indicates the functionality of the transgene and correlates with increased inflammatory marker IL-1β (Fig. 2E). Cytokine ELISA revealed enhanced levels of systemic pro-inflammatory cytokines such as IL1-β, IL-6, and TNF in the serum of Tg-hALOX12B mice compared to the littermate control (Fig. 2F). In addition, Tg-hALOX12B mice manifest increased expression of skin-associated cell adhesion molecules, such as E-selectin, VCAM1, and PECAM, which facilitate the infiltration of CD4 and CD8 cells in the skin (Fig. 2G, H). Furthermore, Tg-hALOX12B also manifested the increased expression of myeloperoxidase (MPO) and collagenase 1 (Col 1), which are well-known markers of skin inflammation-associated pathologies (Fig. 2H). Collectively, these findings suggest a crucial role of *ALOX12B* in the elicitation of psoriasis-like skin inflammatory conditions.

**Figure 2.**
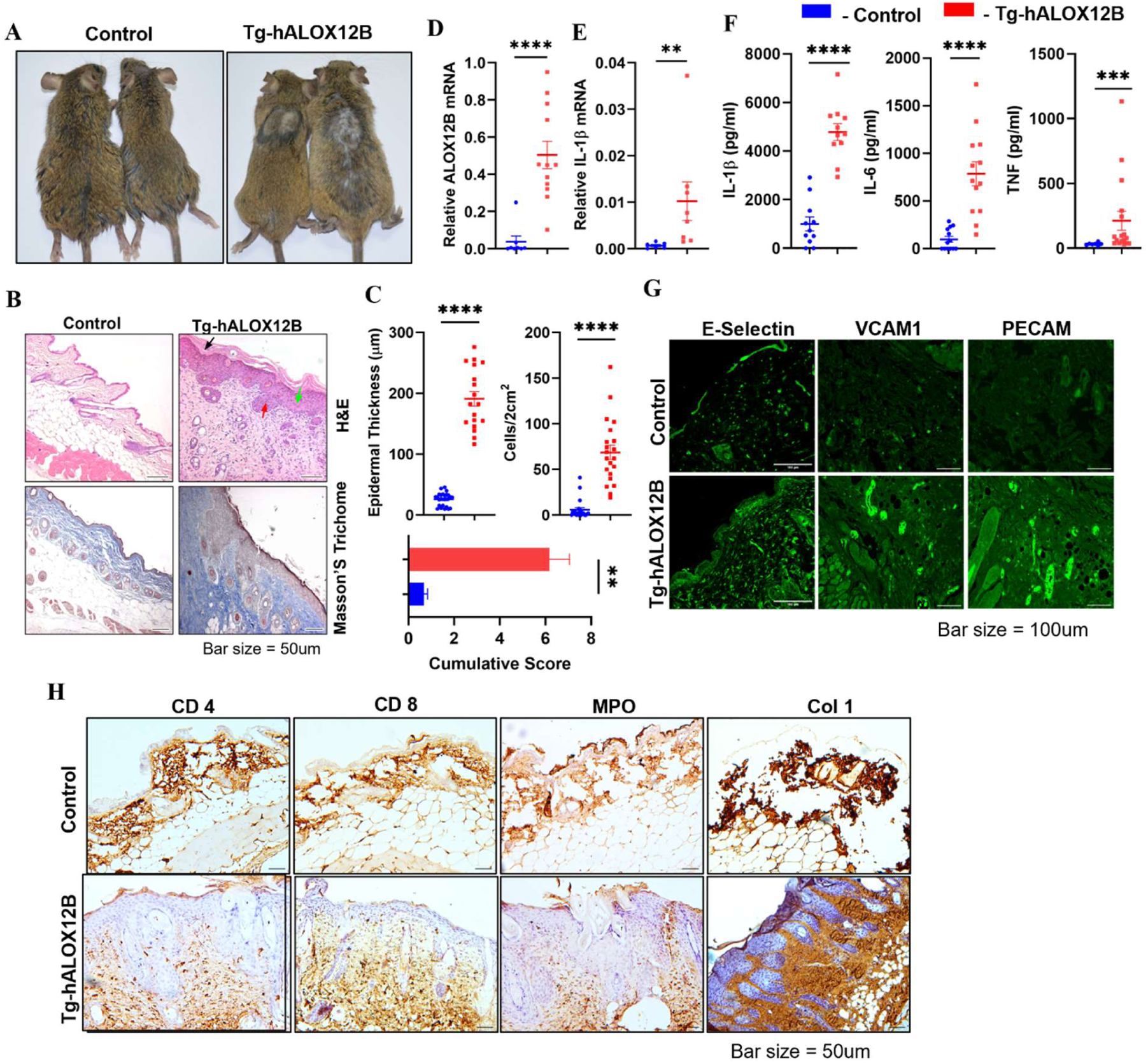
Overexpression of human *ALOX12B* in mice demonstrated a skin abnormality mimicking psoriasis-like skin inflammatory condition. **A)** The humanized transgenic mice (Tg-hALOX12B) phenotypically developed thickened skin patches, and patterned hair loss symptoms closely resembling those observed in skin inflammatory diseases. In contrast, littermate controls did not exhibit these symptoms. **B)** Histological analysis of skin using H&E staining upper panel, showed marked differences between the skin sections of Tg-hALOX12B and littermate control mice. Tg-hALOX12B mice skin histology exhibits several pathological features of skin inflammation; including parakeratosis (green arrow), acanthosis with elongation in the rete ridges (red arrow), and abscess (black arrow). Masson’s trichrome staining lower panel, revealed a significant accumulation of immature collagen (blue color) in the dermis of Tg-hALOX12B mice compared to control mice, a hallmark of skin inflammatory conditions. **C)** Major histopathological parameters were assessed from H&E and Masson’s trichome-stained skin sections of both Tg-hALOX12B and control mice. Tg-hALOX12B mice exhibited a significant increase in epidermal thickness (upper left panel), heightened cellular infiltration in the dermis (upper right panel), and an overall higher combined histopathological score (lower panel), all indicative of skin inflammation. n ≥ 15 vision fields from 5 mice. Scale bars, 50 μM. H&E, hematoxylin, and eosin. **D)** Expression of the human *ALOX12B* gene in the skin of Tg-hALOX12B, in the genotyped animals, as measured by qRT-PCR, *GAPDH* was used as a housekeeping gene control. **E)** Expression of the pro-inflammatory cytokine IL-1β in skin samples of Tg-hALOX12B and littermate controls. **F)** Levels of inflammatory cytokines such as IL-1β, IL-6, and TNF-α detected in the serum of Tg-hALOX12B mice, as estimated by sandwich ELISA. **G)** Dermal expression of key genes associated with psoriasis pathogenesis was evaluated by immunohistochemistry. Tg-hALOX12B mice demonstrated higher expression of cell adhesion molecules, which are known to facilitate leukocyte infiltration into the skin, including E-selectin, VCAM, and PEAM-1 in skin sections, with darker and more prominent spots observed compared to control mice. **H)** Represents the increased infiltration of CD4 and CD8 cells in skin Tg-hALOX12B as compared to control mice supported by cell adhesion molecules. MPO (Myeloperoxidase) and collagenase 1 (col 1) are additional skin inflammation-associated markers. Scale bars, 50 μM Statistical significances were determined by two-tailed unpaired t-test; \**P* ≤ 0.05; \*\**P* ≤ 0.01; \*\*\**P* < 0.001, \*\*\*\**P* < 0.0001. Data are shown as means ± SEM.

### ALOX12B through 12*R*-HETE derives enhanced IL-1β levels by increasing ROS production and inflammasome activation

Having observed that *ALOX12B* is highly expressed in the skin of patients with psoriasis and AD and correlates with clinical inflammatory markers of the disease. Furthermore, the manifestation of inflammation-associated skin pathologies in human *ALOX12B* overexpressing transgenic mice with enhanced levels of pro-inflammatory profile, predominantly IL-1β, suggests the crucial role of *ALOX12B* in inflammatory skin disease pathology. Therefore, we investigated the underlying mechanism by which ALOX12B regulates IL-1β production. ALOX12B mainly converts arachidonic acid into 12*R*-HETE. Previously, it has been reported that 12*R*-HETE concentration is remarkably increased in the epidermis of Psoriatic patients^23,47^. 12*R*-HETE is an active metabolite that binds to several receptors^54,55^ thereby influencing multiple cellular responses^56^. In addition, it has been shown to inhibit Na^+^/K^+^-ATPase function^57^, and its insufficiency in mice triggers heightened pro-inflammatory signals^58^. These observations tempted us to ask the question whether ALOX12B eventually regulates the pro-inflammatory environment in the skin through the enhanced production of 12*R*-HETE. To address this, LPS-primed bone marrow-derived macrophages (BMDM) were stimulated with non-toxic concentrations of purified 12*R*-HETE (Supplementary Data Fig. 7), and cytokine profiles were assessed. Notably, the results showed a significant dose-dependent increase in IL-1β production in the LPS primed 12*R*-HETE treated groups (0.01, 0.1, and 1 µM) compared to that in the LPS or control groups, as measured by ELISA (Fig. 3A). While the levels of IL-6 and TNF were also elevated, their increase was not as marked as that of IL-1β, thereby suggesting that 12*R*-HETE could remarkably influence IL-1β production. Similar results were observed in macrophage cell lines (Supplementary Data Fig. 8). Interestingly, LPS-primed macrophages treated with either *12S*-HETE, *15S*-HETE, or ethanol as a vehicle control were not able to stimulate IL-1β production, highlighting the specificity of 12*R*-HETE in this process (Supplementary Data Fig. 9). Furthermore, to mimic the natural 12*R*-HETE stimulation system, an *in vitro* cell-based ALOX12B overexpression model was established. Murine macrophages were transfected with the human *ALOX12B* overexpressing plasmid construct (pCMV6-hALOX12B) or an empty vector for 18 h using Lipofectamine 3000 followed by LPS priming. Overexpression of *hALOX12B* was confirmed by qRT-PCR (Supplementary Data Fig. 10). The results depicted in Fig. 3B reveal enhanced IL-1β levels in the *ALOX12B* transfected group compared to those in the empty vector control when primed with LPS. Moreover, no change in IL-1β mRNA transcript levels was observed upon 12*R*-HETE treatment alone, indicating the regulation of IL-1β production through ALOX12B/12*R*-HETE at the post-transcriptional level rather than at the transcriptional level (Supplementary Data Fig. 11). IL-1β is a key mediator of various skin inflammatory conditions, including those triggered by infection, environmental factors, and autoimmune diseases^59,60^. The production and maturation of active IL-1β is a complex process that requires two sequential signals: (1) upregulation of IL-1β and NLRP genes, typically induced by TLR ligands, IL-1, or endogenous TNF-α^25^, and (2) assembly of multimeric cytosolic inflammasomes, which are activated by danger signals and subsequently lead to caspase-1 activation and secretion of IL-1β^24,27^.

**Fig 3.**
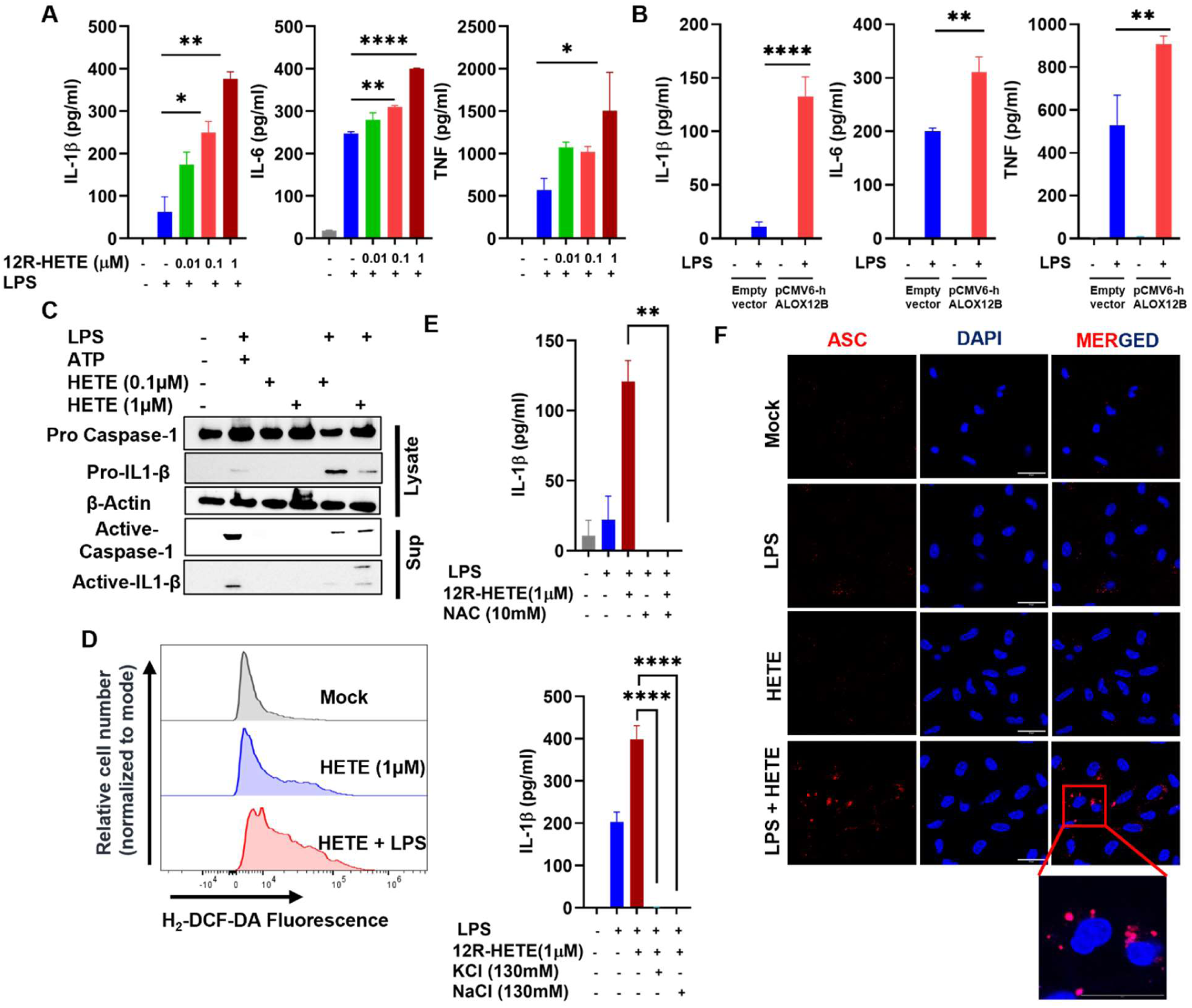
ALOX12B through *12R-*HETE drives enhanced Production of IL-1β by inflammasome activation. **A**) The level of cytokines was assessed from the supernatants of LPS-primed mouse macrophages (500 ng/mL for 5 h) treated with varying concentrations of *12R*-HETE (0.01, 0.1, and 1 µM for 1 h). **B)** Overexpression of human *ALOX12B* (*hALOX12B*) in mouse macrophages by transfecting with *hALOX12B* overexpressing plasmid construct and empty pCMV6 vector control. After 18 h of transfection, cells were primed with LPS for an additional 6 h. The levels of IL-1β, IL-6, and TNF were measured by sandwich ELISA. **C)** Western blot analysis of the supernatant from LPS-primed (500 ng/mL for 5 h), 12R-HETE (0.1 and 1 µM for 1 h), and ATP-treated macrophages. Precursor forms of IL-1β and caspase-1 were detected in the lysates. **D)** ROS levels in LPS-primed (500 ng/mL for 5 h) and *12R*-HETE treated (1 µM for 1 h) macrophages were assessed using H_2_-DCF-DA staining. After 30 minutes of incubation with H_2_-DCF-DA, ROS levels were quantified by flow cytometry. **E)** The involvement of ROS in inflammasome activation was confirmed by pretreatment of LPS-primed macrophages with the ROS inhibitor NAC, 1 h prior to 12R-HETE treatment. Cells were also pretreated with NaCl and KCl, which act as inhibitors of NLRP3 inflammasome activation and the levels of IL-1β assed in the supernatant. NAC treatment resulted in a significantly reduced level of mature IL-1β (upper panel), confirming the role of ROS in inflammasome activation. Similarly, pretreatment with NaCl and KCl also diminished active IL-1β levels, suggesting NLRP3-mediated inflammasome activation through ionic flux modulation (lower panel). **F)** ASC oligomerization, a critical step in inflammasome assembly, was evaluated by immunocytochemistry to visualize ASC speck formation in LPS-primed, 12R-HETE treated macrophages. (left panel: ASC, red; middle panel: DAPI, blue; right panel: merged image; Scale bar size: 20 µM). Statistical significances were determined by One-way ANOVA with Tukey’s multiple comparisons test; ∗, p < 0.05; ∗∗, p < 0.01; ∗∗∗, p < 0.001; ∗∗∗∗, p < 0.0001.

IL-1β and NLRP genes are transcriptionally regulated by the activation of TLRs through various PAMPs, DAMPs, or cellular stress. Among various members of the TLR family, TLR2/1, TLR3, TLR4, and TLR7/8 recognize triacylated lipoproteins, double-stranded RNA, LPS, and single-stranded RNA or free nucleotides respectively^61^. Furthermore, various invading pathogens and cellular stresses are sensed through cytosolic sensors such as NLRs (Nucleotide-binding leucine-rich repeat (LRR) receptors), AIM-2 (Absent in melanoma 2) like receptors, and RIG-1 like receptors^62^. Surprisingly, 12*R*-HETE/*ALOX12B* treated macrophages primed with TLR4 ligand, but not TLR2/1, TLR3, or TLR7/8, induced the release of potent IL-1β (Supplementary Data Fig. 12). Further analysis of the 12*R*-HETE-treated LPS-primed macrophages using western blotting revealed a dose-dependent increase in active caspase-1 and IL-1β, supporting the ELISA data (Fig. 3C). In addition, the transcript level expression of immature IL-1β from 12*R*-HETE treated, LPS primed macrophages was almost comparable with that of LPS-primed macrophages, further confirming that IL-1β release is post-transcriptionally regulated by 12*R*-HETE/ALOX12B but not by transcriptional regulation. These findings prompted us to assess the involvement of cytosolic sensors in the assembly of inflammasome complexes in 12*R*-HETE/*ALOX12B* treated macrophages triggered by the engagement of TLR4 and regulation of the release of IL-1β. NLRP3 is the most widely studied NLRP protein, which responds to a wide range of danger signals and is associated with several inflammation-associated diseases^62^. Although the mechanism through which NLRP3 senses a wide variety of danger signals is not well understood, cellular ROS, K^+^ efflux, and Cl^-^ efflux are known triggers of NLRP3 inflammasome assembly, activation, and IL-1β production^63–65^. Therefore, we examined the production of cellular ROS in 12*R*-HETE-treated macrophages. Interestingly, we found them to be significantly elevated in macrophages treated with 12*R*-HETE, regardless of LPS priming, compared with the mock (Fig. 3D). The levels of active IL-1β were significantly reduced when LPS-primed macrophages were pre-treated with NAC prior to 12*R*-HETE treatment (Fig. 3E, upper panel), indicating that reactive oxygen species (ROS) might be involved in triggering inflammasome assembly^41,65^. To further confirm that the release of IL-1β through 12*R*-HETE/*ALOX12B* is dependent on NLRP3 inflammasome activation, we blocked K^+^ and Cl^-^ efflux by the addition of 130 mM extracellular K^+^ or Cl^-^ ions in LPS-primed macrophages followed by 12*R*-HETE treatment. In agreement with previous results, blockage of K^+^ and Cl^-^ efflux completely blocked the release of potent IL-1β (Fig. 3E, lower panel), suggesting the involvement of NLRP3-mediated inflammasome activation^64^. The levels of release of potent IL-1β from 12*R*-HETE treated, LPS primed macrophages were comparable to those when treated with known danger signals for NLRP3 inflammasome activation, that is, ATP and MSU (Supplementary Data Fig. 13). Finally, we assessed multimeric inflammasome assembly by immunocytochemistry, using ASC protein speck formation as an indicator. The data showed that LPS-primed 12*R*-HETE-treated macrophages exhibited pronounced ASC speck formation (red dots) compared to LPS-only or untreated controls (Fig. 3F). Collectively, these findings suggest that ALOX12B serves as a danger signal by inducing reactive oxygen species (ROS), thereby promoting NLRP3 inflammasome assembly and the subsequent release of active IL-1β.

### Humanized *ALOX12B* overexpressing transgenic mice display enhanced Th17 responses compared to littermate controls

The pathological role of IL-1β in driving the induction of IL-17 producing T helper cells, which contributes to the development of psoriasis and other skin-related disease pathologies, is well established ^66–69^. During skin inflammation, increased production of IL-17 by skin-resident T-cells triggers keratinocyte hyperproliferation, leading to acanthosis, and also acts as a pro-angiogenic factor^70^ in the development of new blood vessels, which exacerbates the inflammatory condition; therefore, IL-17A blocker-based therapies have been lauded for their clinical success in treating Psoriatic Patients^71^. Therapeutics targeting the IL-23 and IL-17 pathways in psoriasis have shown remarkable efficacy. Furthermore, the enhanced production of IL-1β mediated by ALOX12B/12*R*-HETE through inflammasome activation provided us with a unique opportunity to investigate the role of ALOX12B/12*R-*HETE in T cell polarization during *ALOX12B* expression. Our results demonstrated a significant increase in IL-17 producing CD4^+^ T cells in splenocytes of Tg-hALOX12B mice as compared to the littermate controls (Fig. 4A). Furthermore, we also demonstrated increased levels of IL-17 producing CD4^+^ T cells in PBMCs, thereby representing the orchestration of systemic upregulation of inflammatory Th17 cells in Tg-hALOX12B mice compared to the littermates (Fig. 4B). The gating strategy of FACS analysis to segregate IL-17 producing T-cells from skin cells is shown in Supplementary Data Fig. 14.

**Fig 4.**
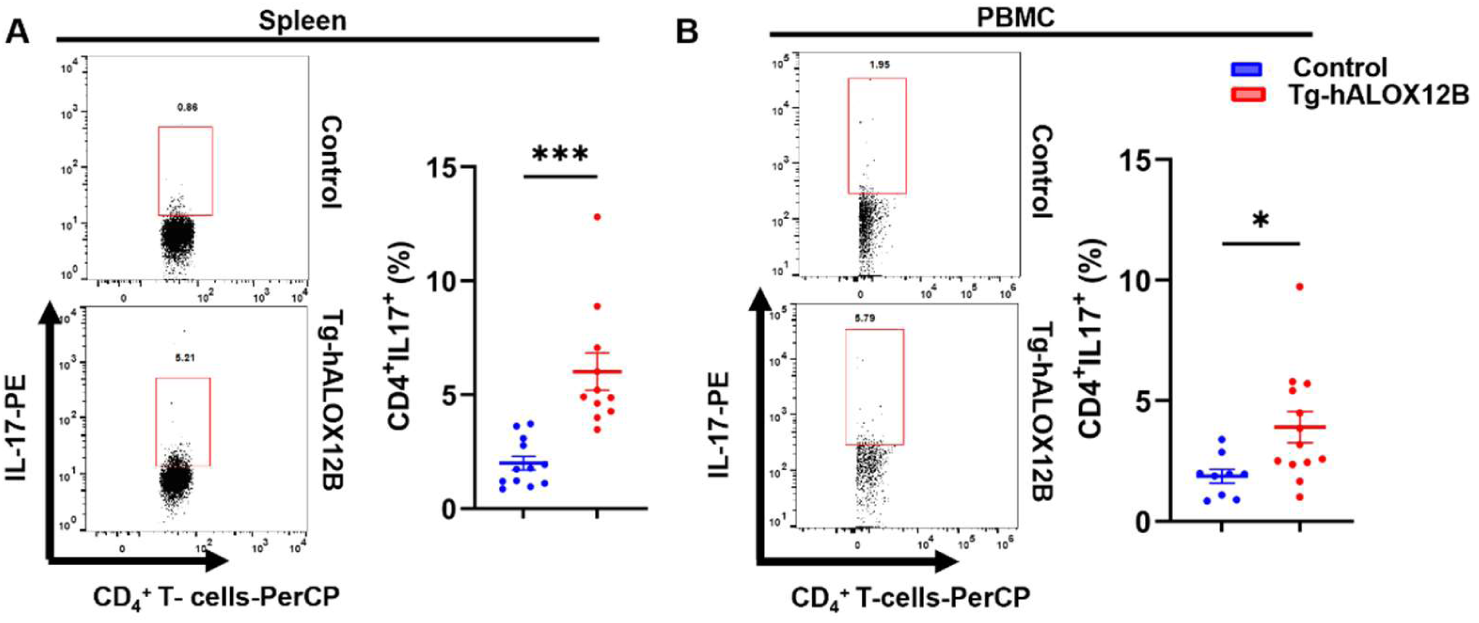
Humanized *ALOX12B* overexpressing transgenic mice display enhanced Th17 responses compared to littermate controls. **A)** Effect of *ALOX12B* in modulation of IL-17 producing T-cells in splenocytes, and **B)** PBMCs were observed in the Tg-hALOX12B mice compared to littermate control mice. The FACS plots are representative of three or more than three independent experiments. Each dot in the bar graph adjacent to the respective FACS plots is a representative of an individual animal. Statistical significance between Tg-ALOX12B and control groups was determined using Student’s t-test, with ∗, p < 0.05; ∗∗, p < 0.01; ∗∗∗, p < 0.001.

### Rationally designed novel ALOX12B inhibitor alleviates skin inflammatory symptoms in *ALOX12B* overexpressing transgenic mice

Given the involvement of ALOX12B in regulating skin inflammation, the design of a specific ALOX12B inhibitor is a promising strategy for developing novel and effective therapeutic interventions for inflammatory skin diseases. However, the development of such inhibitors presents a significant challenge because of the scarcity of known ALOX12B inhibitors. Additionally, the lack of a three-dimensional crystal structure of ALOX12B in complex with its substrate has hindered efforts in medicinal chemistry, leaving this area largely unexplored ^72^. Nonetheless, given the high degree of structural similarity among the members of the LOX gene family^73,74^ particularly in the active site region, baicalein^75–77^ a known non-selective lipoxygenase inhibitor, has emerged as a potential scaffold for the development of novel ALOX12B inhibitors. Consequently, by optimizing the structure of baicalein, we designed a new flavone-based scaffold (Scaffold-**I**) for the present study (Fig. 5A). Subsequently, *in silico* screening of various analogs derived from Scaffold-**I** was performed to assess the potential of the designed flavone derivatives. A set of promising compounds was selected based on their optimal binding scores within the enzyme’s substrate-binding pocket and compared to the binding score of baicalein (Table S3) for further synthesis. Given that the crystal structure of human ALOX12B is not yet available, homology modelling was employed to generate a model of the enzyme (Supplementary Data Fig. 15), which was used for docking studies. The targeted compounds obtained from the *in silico* analysis (Table S4) were synthesized according to the reaction scheme outlined in Fig. 5B and Scheme SB, C, and D (Supplementary Methods), and their characterization was performed using spectral data (Supplementary Data Fig. 16). Subsequent screening using an *in vitro* hALOX12B inhibition assay revealed that a few compounds exhibited encouraging inhibitory activities against hALOX12B (Table S4). Notably, compound **6a** showed promising inhibition of hALOX12B, and its *in silico* binding affinity (Fig. 5C) correlated well with the *in vitro* inhibition results (Table S5a). Furthermore, in LOX selectivity studies of compound **6a**, selective inhibition of hALOX12B over hALOX12, hALOX15, and hALOX15B was observed (Table S5b).

**Fig 5.**
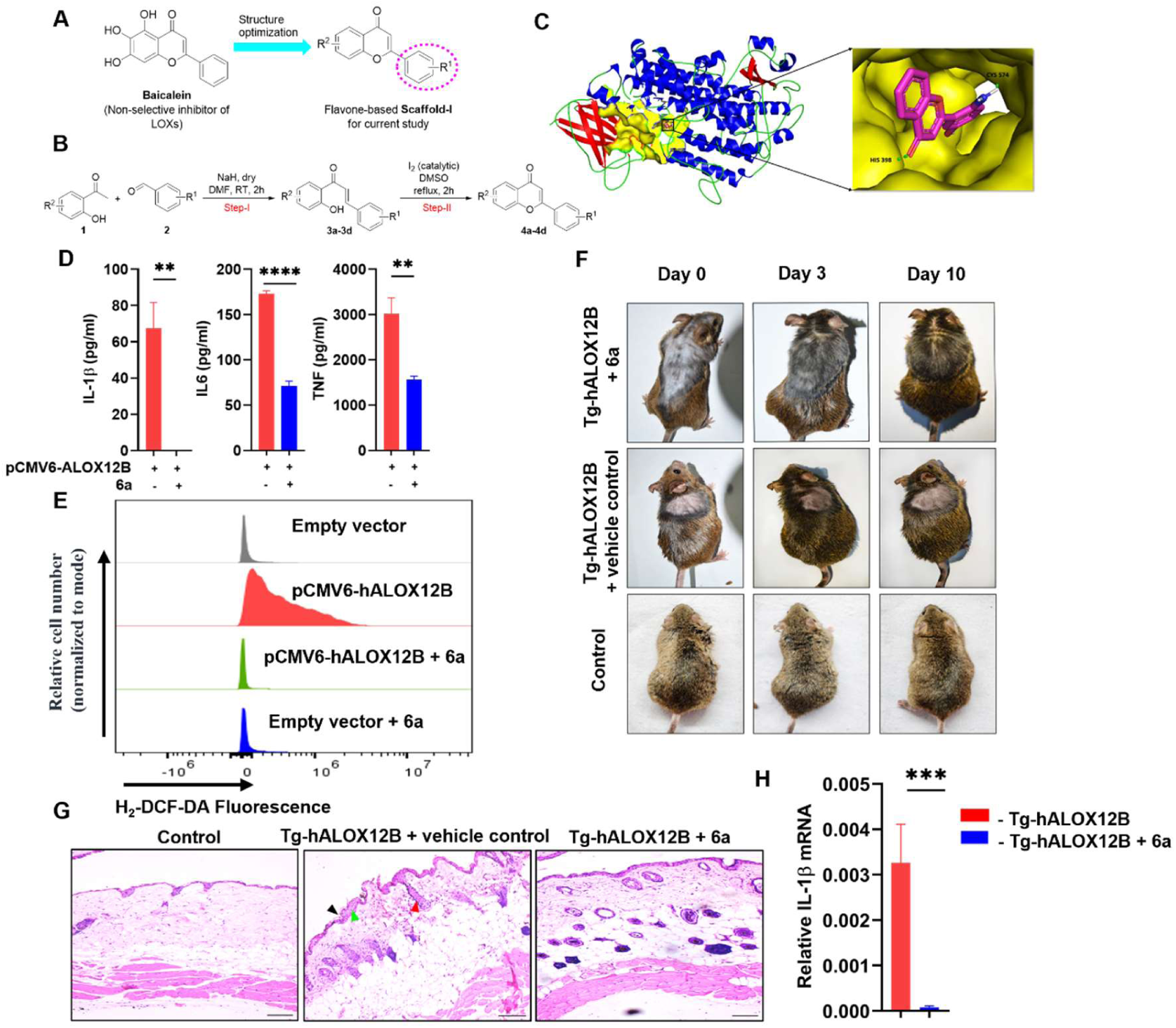
Rationally designed novel ALOX12B inhibitors mitigate skin inflammatory symptoms and associated molecular markers in transgenic mice. **A)** Depicts the design of potential hALOX12B inhibitors. The flavone-based Scaffold-I was developed through the structural optimization of the non-specific LOX inhibitor, baicalein. Various compounds were initially designed by modifying the R^1^ group in Scaffold-I and then screened *in-silico* against the ALOX12B enzyme. A selection of molecules, based on binding affinity analysis, was chosen for synthesis. **B)** A general chemical synthetic scheme outlining the synthesis of the flavone backbone based on Scaffold-I. **C)** The human ALOX12B protein model, generated via homology modeling using AlphaFold software, is shown. The binding mode of the hALOX12B inhibitor 6a, which demonstrated encouraging *in-vitro* inhibition within the series, is shown in the binding pocket of hALOX12B. **D)** The inhibition potential of compound **6a** was assessed *in-vitro* at a concentration of 10 µM by measuring the levels of mature IL-1β production along with TNF and IL-6 in LPS-primed macrophages, overexpressing pCMV6-hALOX12B as compared to the vector control. **E)** Intracellular ROS levels were analysed in cells transfected with pCMV6-hALOX12B, treated with or without compound 6a. **F)** For the *in-vivo* studies, compound **6a** was homogenized in petroleum jelly base and applied daily for 10 days on affected skin patches of Tg-hALOX12B mice at 50 mg/kg body weight. After 10 days, animals were sacrificed for downstream analysis. **G)** H&E staining was done to assess the skin inflammatory features; parakeratosis (green arrow), acanthosis with elongation in the rete ridges (red arrow) and abscess (black arrow) in compound **6a** treated animals compared to the vehicle control. **H)** The expression of IL-1β in the skin was assessed by qRT-PCR. Data are representative of three animals (n = 3), and statistical significance was calculated using Student’s t-test, with p < 0.05 considered significant. Scale bars, 100 μM. Statistical significances were determined by One-way ANOVA with Tukey’s multiple comparisons test; ∗, p < 0.05; ∗∗, p < 0.01; ∗∗∗, p < 0.001; ∗∗∗∗, p < 0.0001.

To further evaluate the anti-inflammatory effects of compound **6a** *in vitro*, we treated *ALOX12B*-overexpressing LPS-primed macrophages with compound **6a** at a concentration of 10 µM (based on the MTT assay, Supplementary Data Fig. 17). We observed a marked inhibition of active IL-1β production by compound **6a** when compared to *ALOX12B* overexpressing LPS-primed macrophages (Fig. 5D). The levels of IL-6 and TNF were also decreased, although not to the same extent as IL-1β, which showed a substantial reduction. Notably, compound **6a** also inhibited ROS production in ALOX12B-overexpressing LPS-primed macrophages compared to the empty vector control (Fig. 5E). These robust findings underscore the potential of compound **6a** as a novel and selective ALOX12B inhibitor and a potential lead compound for the development of therapeutic regime for the treatment of inflammatory skin disease. Hence, these findings prompted us to study the preclinical, safety, toxicity, and *in vivo* profiles of compound **6a.**

### Pharmacokinetic and toxicity profiling of 6a

To evaluate the drug-like properties of compound **6a**, preliminary pharmacokinetic (PK) and toxicity studies were performed. An *in vitro* metabolic stability study of **6a** at a test concentration of 5 μM in mouse liver microsomes demonstrated favorable stability, with a half-life (t_1/2_) of 495 minutes and an intrinsic clearance of 5.6 μL/min/mg protein (Table S6, Supplementary Data Fig. 18). However, following intravenous (IV) administration of **6a** at a dose of 10 mg/kg body weight, the compound exhibited a non-optimal PK profile in mice (Supplementary Data Fig. 19). Subsequently, a single-dose dermal PK study of **6a** (50 mg/kg body weight) was performed in mice to assess its dermal PK properties and evaluate its suitability for topical applications (Table S7). As shown in Tables S8 and S9 and illustrated in Supplementary Data Fig. 20, no clinical signs or mortality were observed in the experimental animals, and the peak tissue concentration was observed 4 h after post-application, with the highest tissue concentration (C_max_) reaching 71,766.33 ng/g. Importantly, the optimal tissue concentration of **6a** was maintained for over 24 h (Supplementary Data Fig. 20a) in the experimental animals. In contrast, a marginal peak plasma concentration (C_max_ = 28.79 ng/mL) was observed in the mice (Supplementary Data Fig. 20b). These findings suggest that compound **6a** possesses favorable dermal PK properties, making it a promising candidate for topical applications. Moreover, given the cutaneous nature of inflammatory skin conditions, topical administration of **6a** may offer enhanced and targeted therapeutic efficacy, while minimizing the adverse effects typically associated with systemic delivery.

Next, an acute dermal toxicity study of compound **6a** was conducted in Wistar rats using the fixed-dose procedure to assess its dermal toxicity. The study began with a dose-range-finding phase, followed by the main study at a dose level of 2000 mg/kg body weight. Throughout the 14-day observation period, the animals did not exhibit clinical signs of toxicity or mortality. Additionally, gross pathological examination at the end of the observation period revealed no macroscopic lesions. Based on these results, the LD_50_ of **6a** was determined to be > 2000 mg/kg body weight under the tested conditions (Table S10). Thus, according to the Globally Harmonized Classification System (GHS) for Chemical Substances and Mixtures, **6a** was classified as “Category 5” (Table S11). Additionally, the potential cardiac toxicity of compound **6a** was assessed by evaluating its ability to block hERG (IKr) ion channels *in vitro* using the whole-cell patch-clamp technique. At test concentrations ranging from 0.3 to 30 µM, compound **6a** did not cause a significant reduction in the hERG current (9-14 %) (Table S12), indicating that it is unlikely to pose a risk of cardiac toxicity.

### Therapeutic efficacy of 6a in Tg-hALOX12B mice

Based on the favorable dermal PK and safety characteristics, we evaluated the efficacy of compound **6a** in hALOX12B overexpressing transgenic mice by administering its local formulation at a dose of 50 mg/kg for 10 days. The treatment resulted in a remarkable improvement in symptoms over the study period compared to the vehicle control (Fig. 5F). Histopathological analysis of the treated skin showed normalized epidermal thickness and a reduction in inflammatory features such as parakeratosis (green arrow), acanthosis with elongation of the rete ridges (red arrow), and abscess formation (black arrow) (Fig. 5G). Additionally, we investigated inflammatory gene signatures, revealing reduced expression of IL-1β in the skin of animals treated with compound **6a** compared to that in the untreated groups (Fig. 5H).

Overall, our identified hALOX12B inhibitor, **6a**, not only downregulated the levels of IL-1β, IL-6, and TNF *in vitro* but also inhibited the expression of IL-1β *in vivo*. These results demonstrate the critical role of ALOX12B mediated IL-1β production and its involvement in inflammatory T cell polarization in the skin. Therefore, targeting the ALOX12B/IL-1β axis with a hALOX12B inhibitor, such as compound **6a** could provide a novel strategy for the potential management of skin inflammatory diseases such as psoriasis and atopic dermatitis (AD).

## Discussion

Despite the availability of a wide variety of information regarding skin inflammation, yet there is a dearth of knowledge regarding the complexity of molecular changes responsible for immune dysregulation and disease pathology. Therefore, a global understanding of the inflammatory correlates in the skin is necessary to gain insights into both health and disease conditions. To delineate the global architecture of skin inflammation, we adopted gene expression meta-analysis on the publicly available datasets in the skin biopsies of Psoriatic and Atopic dermatitis patients and further confirmed the gene meta-profile in the skin tissues of the Psoriatic patients reporting to well-recognized hospitals in Hyderabad. Our data revealed a wide variety of genes upregulated in the Psoriatic and AD patients. However, we identified a gene signature comprising *ALOX12B*, which was significantly upregulated across all the patients’ samples and positively correlated with key inflammatory markers such as IL-1β, IL-17, and IL-23. ALOX12B is an enzyme that converts arachidonic acid into *12R*-HETE^13^. Several seminal studies described the crucial role of *ALOX12B* in the development of skin. For instance, mutations in the coding regions of *ALOX12B* have been linked to autosomal recessive congenital ichthyosis (ARCI), a rare genetic skin disorder, characterized by impaired epidermal barrier function, erytheroderma, and hyperkeratosis^16–18^. Moreover, *ALOX12B* deficient mice exhibit perinatal lethality due to severe defects in skin barrier formation^19^, underscoring the enzyme’s critical role in the proper development of skin. Further, skin-enriched psoriatic DEGs” (SEPDs) identified in our study predominantly involved *ALOX12B*, along with *LAMB4*, *IL37*, and *CST6*. Notably, *LAMB4*, *IL37*, and *CST6* were consistently downregulated across most of the psoriatic skin datasets, while *ALOX12B* was significantly upregulated. This skin-specific expression pattern of *ALOX12B*, combined with its correlation with inflammatory mediators, suggests its potential role in driving skin inflammation. Furthermore, *ALOX12B* appears to be primarily expressed in the skin^44^, unlike other ALOX family members are more widely distributed across tissues^78^.In addition, accumulation of *ALOX12B’s* enzymatic product, 12*R-*HETE in the skin, has been reported in psoriasis and other inflammatory skin diseases^20–23^, further supporting its involvement in skin inflammation and disease pathogenesis. In line with these findings, our clinical validation data in the psoriatic patient skin samples reveal upregulation of *ALOX12B* and other known DEGs, notably IL-1β, IL-36G, and IL-22, as well as genes associated with skin tissue organization, including *KYNU, S100A12, and HPSE.* These results are consistent with the established roles of these genes in the pathogenesis of inflammatory skin diseases^79,80^, further suggesting that *ALOX12B* might function as a key molecular switch in triggering skin inflammation and associated pathologies.

To elucidate the pathological role of *ALOX12B*, we developed a humanized transgenic mouse model expressing the human ALOX12B enzyme in the skin (Tg-hALOX12B). Notably, Tg-hALOX12B transgenic mice exhibited skin abnormalities resembling chronic inflammatory conditions, including epidermal hyperplasia, immune cell infiltration, and dermal fibrosis, which are hallmark features of several inflammatory skin conditions^6,7^. Histopathological analysis of the skin further confirmed these abnormalities, suggesting that overexpression of *ALOX12B* in the skin can induce inflammation and pathological changes in the skin akin to psoriasis-like symptoms.

Given the heightened levels of IL-1β observed in both psoriatic patients’ skin and the skin and serum of Tg-hALOX12B, we further explored how *ALOX12B* regulates IL-1β production. Our findings demonstrated that 12*R*-HETE triggered a dose-dependent increase in IL-1β production in LPS-primed macrophages without any changes at the transcript level. This suggests that ALOX12B may be involved in the post-translational regulation of IL-1β as it requires two signals: first, to regulate the transcription of the gene known as priming signals, and second, as a danger signal for triggering inflammasome activation^81^. We found that inhibiting reactive oxygen species (ROS) or blocking ion efflux dampened the IL-1β response, indicating that *12R*-HETE triggers NLRP3 inflammasome assembly, with ROS being a major activator, as ROS and ion efflux have been reported to be linked with NLRP3 inflammasome activation^82^. Furthermore, ASC oligomerization by ASC speck formation was observed in LPS-primed *12R*-HETE-treated macrophages, reinforcing the idea that *ALOX12B* plays a pivotal role in inflammasome activation. In brief, *ALOX12B* enhances mature IL-1β levels by triggering inflammasome assembly through the ROS-NLRP3-ASC-Caspase-1 axis.

The significance of IL-1β in inflammatory skin diseases, particularly in the differentiation of Th17 and γδT17 cells, is well-established^59,68,83^. In human *ALOX12B* overexpressing transgenic mice, we observed an expansion of IL-17-producing T cells, which was regulated by the levels of active IL-1β, demonstrating the critical role of *ALOX12B* in driving IL-17 responses^66^.

Given the involvement of *ALOX12B* in skin inflammation, we designed a specific ALOX12B inhibitor, compound **6a**, based on the structure of biaclein, a known non-selective lipoxygenase inhibitor, which showed significant efficacy in both *in vitro* and *in vivo* models. Topical application of compound **6a** to Tg-hALOX12B transgenic mice alleviated inflammatory symptoms, normalized skin morphology, and reduced key inflammatory markers such as IL-1β. The acceptable preclinical parameters, such as topical pharmacokinetics along with acute dermal and cardiac safety profiles, suggest that compound **6a** has potential as a prospective agent for treating inflammatory skin diseases.

In summary, our study highlights the critical role of *ALOX12B* in the pathogenesis of inflammatory skin conditions, particularly through the ROS production, regulation of inflammasome activation, and IL-1β/Th17 axis. Pharmacological targeting of ALOX12B with a novel inhibitor such as compound **6a** offers a promising therapeutic strategy for treating skin inflammation. However, further studies are warranted to fully understand the mechanisms underlying *ALOX12B* contribution to skin inflammatory diseases in the human settings and to optimize the clinical translation of ALOX12B-targeted therapies.

## Acknowledgments

We are thankful to Dr. Elian Fusch from Rockefeller University, USA for providing us pGE3Z.K14 construct. We also acknowledge Dr. Satish Kumar (Scientist, CCMB) and Mrs. B Jyothi Lakshmi. (Sr. Technical Officer, Animal House, CCMB) for their support in the generation of Tg-hALOX12B mice. We are thankful to the School of Life Sciences for providing access to Confocal Microscopy and Flow cytometry facilities.

## Funding

We are grateful for the financial support provided by the DST-DPRP, New Delhi, India [VI-D&P/560/2016-17/TDT (G-1)], Institute of Eminence (IoE) (UoH/IoE/IIRC-23-003) Institute of Eminence (IoE) (UoH/IoE/RC1/RC1-20-017), Indian Council of Medical Research (ICMR) (34/11/2019 T/F/NANO/BMS), ICMR Extra Mural Small Grant (EMDR/SG/12/2023-5626), and Department of Biotechnology—Biotechnology Industry Research Assistance Council (DBT-BIRAC) (DBT/04/0401/2019/01546). S.S. is supported by the DBT-SRF Fellowship from the Department of Biotechnology (DBT/2022-23/UOH/1985), and F. A. is supported by the DHR-Young Scientist Scheme (R.12014/63/2020-HR). We also thank the DBT-builder grant from the School of Life Sciences (BT/INF/22/SP41176/2020).

## Data Availability Statement

All relevant datasets generated or analyzed during the study are included in the manuscript and its supplementary materials.

## Ethics declarations

All animal experiments were conducted in accordance with the guidelines and regulations approved by the Institutional Animal Ethics Committee (IAEC) of [University of Hyderabad; UH/IAEC/NK/2023-1/43/R1], in compliance with the Committee for the Purpose of Control and Supervision of Experiments on Animals (CPCSEA; 151/GO/ReBi/S/99/CPSCEA), Government of India.

Human samples and data were collected following protocols approved by the Institutional Ethical Committee, University of Hyderabad; UH/IEC/2019/210. Informed written consent was obtained from all participants prior to sample collection.

## Competing interests

The authors declare no competing interests.

## Contributions

Conceptualization: S.S., F.A., N.K. methodology: S.S., F.A., N.F., C.S.N., H.B., R.A.K., S.A., S.B., K.K.R., M.J., S.N., N.S., A.M.P., T.P., V.V., M.J.M.K. investigation: S.S., F.A., N.K. data visualization and compilation of figures: S.S., C.S.N., N.K. computational analysis: C.S.N. funding acquisition: S.O., M.P., P.R., N.K. project administration: N.K. study supervision: N.K. writing-original draft: S.S., F.A., N.K. writing-review & editing: S.S., F.A., C.S.N., H.B., M.P., N.K.

## Notes

### Competing Interest Statement

The authors have declared no competing interest.

