## supplementary materials for "ALOX12B overexpression in the skin drives inflammasome/Th17 signaling axis to promote inflammation in the mouse model and human patients"

### Supplementary methods

#### Schematic representation of the chemical synthesis of novel ALOX12B inhibitors

**Scheme SA:**

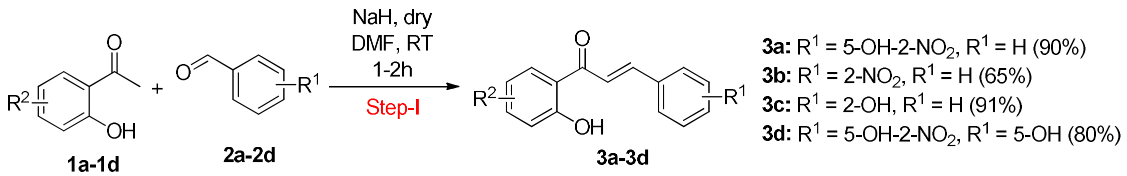


#### General procedure for the synthesis of intermediate 3 (3a-3d)

Sodium hydride (3 mmol) was added slowly to an ice-cold solution of appropriate acetophenone (**1**, 1 mmol) in dry DMF (20 mL) under a nitrogen atmosphere, and the reaction mixture was stirred at ambient temperature for 20 min. Subsequently, 2-nitrobenzaldehyde (**2**, 1.1 mmol) was added to the reaction mixture, and the solution was stirred at ambient temperature for 1-2 h. The disappearance of starting materials was monitored by TLC. After completion of the reaction, the mixture was poured into ice-cold water, and the pH was adjusted to ~6 using an aqueous HCl solution. The obtained solid was removed by filtration and washed with warm water (2 × 50 mL) and *n*-hexane (2 × 50 mL) to give the intermediate **3**.

**Scheme SB:**

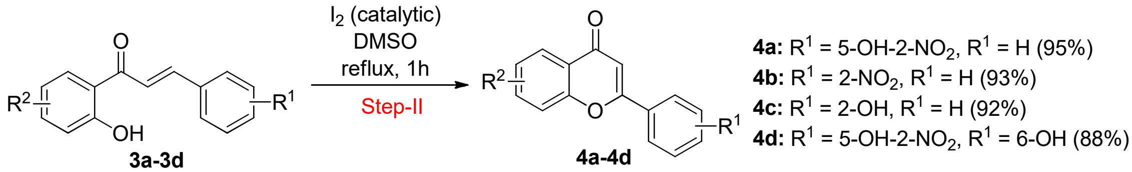


#### General procedure for the synthesis of flavones 4 (4a-4d) from the chalcone intermediates (3a-3d)

Intermediate **3** (1 mmol) and a catalytic amount of $I_2$ were dissolved in DMSO (10 mL), and the solution was refluxed (the temperature of the heating oil bath was maintained at 200 °C) for 1h. After completion of the reaction (monitored by TLC), the reaction mixture was cooled to room temperature and diluted using cold water (50 mL). The mixture was extracted with EtOAc (3 × 30 mL). The organic layers were collected, combined, washed with water (3 × 20 mL), dried over anhydrous sodium sulfate, filtered, and concentrated under vacuum. The residue obtained was purified by column chromatography using 40 % of ethyl acetate/*n*-hexane as the eluent to obtain the desired compound **4**.

**Scheme SC:**

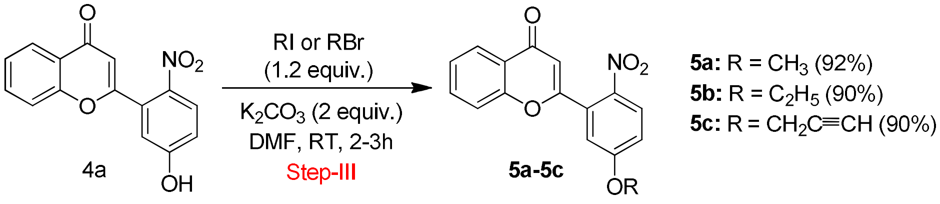


**General procedure for the synthesis of flavones 5 (5a-5c) from the flavone 4a**

To a solution of compound **4a** (0.283 g, 1 mmol) in DMF (10 mL), RI or R'Br (1.2 mmol) and $K_2CO_3$ (0.28 g, 2 mmol) were added, and the reaction mixture was stirred for 2-3 h at room temperature. After the disappearance of the starting material (monitored by TLC), ice-cold water (50 mL) was added, and the reaction mixture was extracted with EtOAc (2 × 50 mL). The organic layers were then collected, combined, dried over anhydrous $Na_2SO_4$ and evaporated to obtain the crude product. The crude product was then purified by using column chromatography with 30 % EtOAc/*n*-hexane as the eluent.

**Scheme SD:**

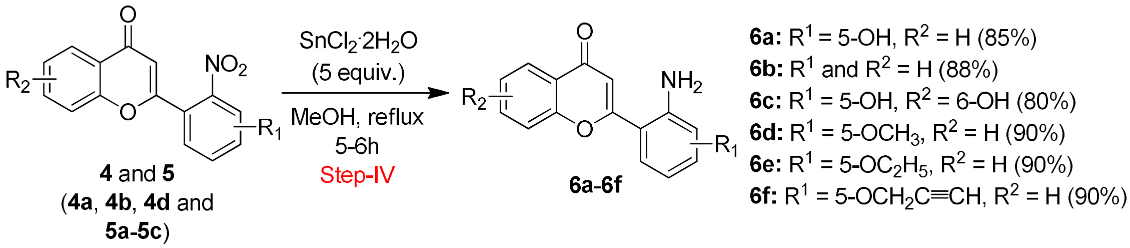


**General procedure for the synthesis of flavones 6 (6a-6f) from flavone 4 and 5**

SnCl2 · 2H2O (5 mmol) was added to a stirring solution of compound **4** or **5** (1 mmol) in MeOH, and the resultant reaction mixture was refluxed until the starting material disappeared (monitored by TLC). Then, ice-cold water was added to the reaction mixture, and the residue obtained was diluted with 2N NaOH (50 mL). The resultant reaction mixture was then extracted with EtOAc (3 × 50 mL), and the organic layers were collected, combined, dried over $Na_2SO_4$, filtered, and concentrated to afford the crude product. The crude product was then purified by column chromatography using 40 % EtOAc/*n*-hexane as eluent.

**Supplementary method for metabolic stability of 6a in mouse liver microsomes**

Test item **6a** or control item diclofenac sodium or imipramine hydrochloride was incubated at a concentration of 5 μM with mouse liver microsomes (0.25 mg/mL) in the presence or absence of cofactors (NADPH regeneration system). A 100 μL reaction mixture was removed at specified time points (with cofactors: 0, 15, 30, 60, and 120 min; without cofactors: 0 and 120 min), and the reaction was stopped by the addition of stop solution. The samples were extracted in the presence of an internal standard and analyzed using LC-MS/MS. The amount of test/control items remaining at a specified incubation period was calculated by comparing the peak area ratio at each time point with the peak area ratio at 0 min.

**Supplementary method for single Dose Dermal Pharmacokinetic Study of 6a in Mice**

The application site was prepared by clipping the fur from the lumbar region using a clipper approximately 24 h before application. Care was taken to avoid skin abrasion. Only animals with healthy, intact skin were used. To the weighed test item of 40 mg, 4 mL DMSO was added and vortexed to solubilize the test item. The test items were evenly applied to the clipped skin area on the dorsal side of the experimental animal based on body weight (Table S7). At specified periods, blood and skin were collected at (predose, 1, 2, 3, 4, 6, and 24 hours) after application of 50 mg/kg of the formulation, and the animals were sacrificed by decapitation. A skin incision was made (application site), separated, and washed with normal saline. The removed skin was weighed and transferred into a tube, and homogenized by adding sufficient buffer. The matrix concentration was determined using the fit for the LC/MSMS method. Post-application of the test formulation to mice, maximum blood samples were collected from 3 mice at each pre-defined time point (predose, 15 min, 30 min, 2 h, 12 h, 24 h, days 2, 3, 4, 5, 6, 7, 8, 15, 21) under isoflurane anesthesia into pre-labelled tubes containing an anticoagulant. The tubes were then centrifuged at 14,000 rpm for 5 min at 4 °C to harvest plasma samples and stored at − 80 ± 10°C until further analysis. Following single dermal application of **6a** to mice, the time for peak tissue concentration ($T_{max}$) was 4.00 h post-dose at 50 mg/kg/day. The plasma and tissue samples were quantified with **6a** till 1.00–24.00 h post dose ($T_{last}$).

**Supplementary method for intravenous pharmacokinetic (IV) profile of 6a**

Male BALB/c Mice aged 8-9 weeks were used for experimentation after a minimum 3 days of acclimatization. Non-fasted animals were administered with compound **6a** in a recommended vehicle (5% v/v DMA + 5% v/v Tween 80 + 10% v/v Solutol HS 15 + 80% v/v Phosphate buffer,10 mM, pH 8.0) by intravenous route with a dose of 10 mg/kg body weight and at a dose volume of 10 mL/kg body weight. The blood specimens were collected from retro-orbital plexus up to 24 h post-dose. Collected blood specimens were centrifuged at 6000 rpm, 4 ºC for

10 minutes, and plasma was separated and stored at -80ºC until analysis. Plasma concentrations of **6a** were quantified by LC-MS/MS, and the plasma pharmacokinetic parameters were determined by non-compartmental analysis using Phoenix WinNonlin 6.3 software.

**Supplementary method for Acute dermal toxicity study of 6a in Wistar rats**

The "Acute Dermal Toxicity" study was conducted as per Organization for Economic Co-operation and Development Guideline for Testing of Chemicals "Acute Dermal Toxicity: Fixed Dose Procedure" (Section-4, No. 402; adopted 9th October 2017). Initially, the study was performed as a dose-range-finding study using one female animal at a dose of 2000 mg/kg body weight. No clinical signs or mortalities were observed. Therefore, the main study was conducted using two additional female animals at a dose level of 2000 mg/kg body weight. On test day 0, the quantity of test item **6a** was calculated for each animal based on its body weight and the equivalent weight of the test item. The test item was applied to the clipped area of the skin of each animal with a porous gauze dressing, bandaged with non-irritating adhesive tape, and kept in contact throughout the 24 h exposure period. The test site was further covered with an elastic adhesive bandage to retain gauze dressings. After a 24 h exposure, the dressings were removed, the skin was gently wiped with cotton soaked in RO water, and skin reactions were assessed.

**Supplementary method for *In-vitro* hERG channel (ikr) assay**

The NPC-1 chip of the Port-a-Patch was filled with 5µl of Internal buffer solution (KCl 50 mM, NaCl 10 mM, KFl 60 mM, EGTA 20 mM, Hepes/KOH 10 mM, pH 7.2 ± 0.05, osmolarity: 285 ± 5 mOsmol) and screwed onto the chip holder. The Faraday cage was then fixed on the chip holder, such that the external electrode was placed near the chip aperture. 5µl of External buffer solution (NaCl 140 mM, KCl 4 mM, MgCl2 1 mM, CaCl2 2 mM, D-Glucose monohydrate 5 mM, Hepes/NaOH 10 mM, pH 7.4 ± 0.05, osmolarity:298 ± 5 mOsmol) was added to the center of the aperture so that the external electrode was in contact with the external buffer solution. The experiment was initiated. Once the set threshold of resistance is attained (i.e., 2-3.5 MOhm), 5 µl of cell suspension was added into the middle of the external buffer solution droplet after the Suction Control unit has generated the suction pulse. Suction automatically pulls a cell to the aperture, resulting in an increase in the chip resistance. When the cell is caught and the set threshold for the resistance reaches (i.e., 5 MOhm), the software recognizes this increase and proceeds to the next sealing step. 20 µl of seal enhancer solution (NaCl 80 mM, KCl 3 mM, MgCl2 10 mM, CaCl2 35 mM, HEPES (Na+salt)/HCl 10 mM pH

7.4 ± 0.05, osmolarity: 298 ± 5 mOsmol) was added after 2-3 washes. When the threshold resistance was reached, PatchControl automatically moved to the step of improving the seal. Once the desirable resistance (Rpip) was reached, the whole-cell configuration was attained and maintained. The voltage protocol consisted of depolarization of the cell membrane to + 40 mV for 500 ms (for channel activation) from a holding potential of − 80 mV. Cells were stimulated every 10 seconds using this protocol at each drug concentration. 6a was tested at 0.1, 0.3, 1, 3, 10, and 30 μM (0.1 % v/v DMSO) concentrations.

**Supplementary results**

**a)**

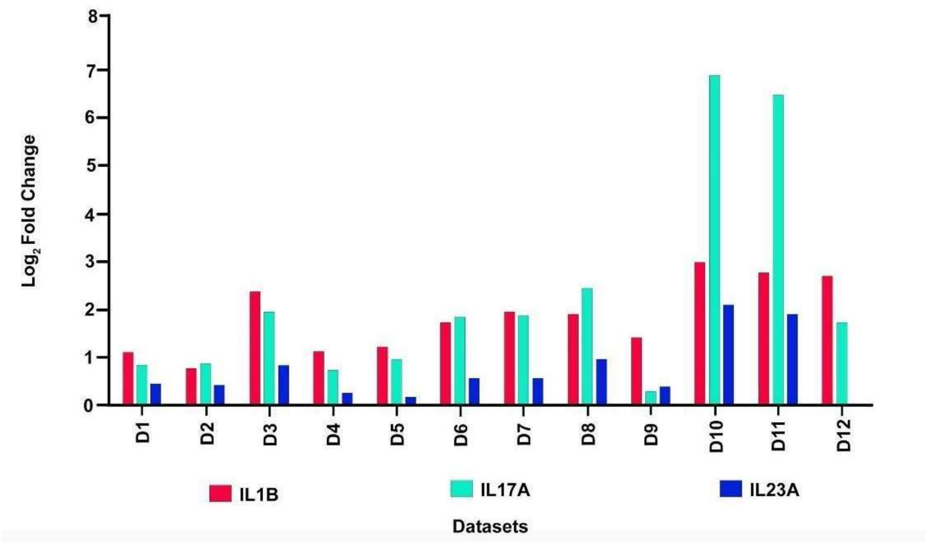

**b)**

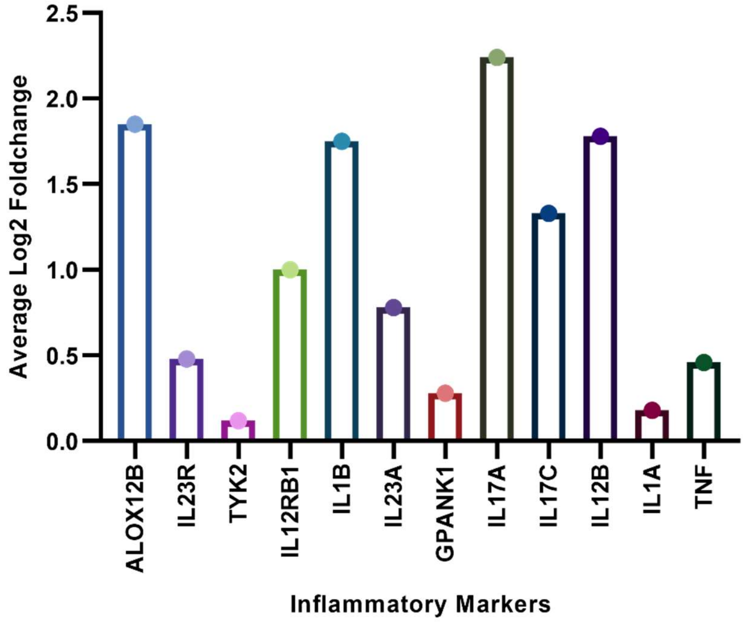


**Supplementary Data Figure 1: Differential expression of key inflammatory markers in psoriasis. (a) Dataset-Specific $log_2$ fold change of selected inflammatory cytokines:** $Log_2$ fold change values for *IL1β*, *IL17A*, and *IL23A* gene expression were determined across 12 independent datasets derived from psoriatic patient skin biopsies. Consistent upregulation of *IL1β*, *IL17A*, and *IL23A* was observed in the differential expression analysis of each dataset, indicative of their robust involvement in the psoriatic inflammatory response. **(b) Average $log_2$ fold change of inflammatory markers:** A bar plot was generated to visualize the average $log_2$ fold change for a panel of key inflammatory markers, including *IL1β*, *IL17A*, *IL23A*, and *ALOX12B*. This analysis revealed a concordant pattern of upregulation for most assessed inflammatory markers alongside *ALOX12B*, suggesting a coordinated increase in pro-inflammatory gene expression within psoriatic lesions.

a)

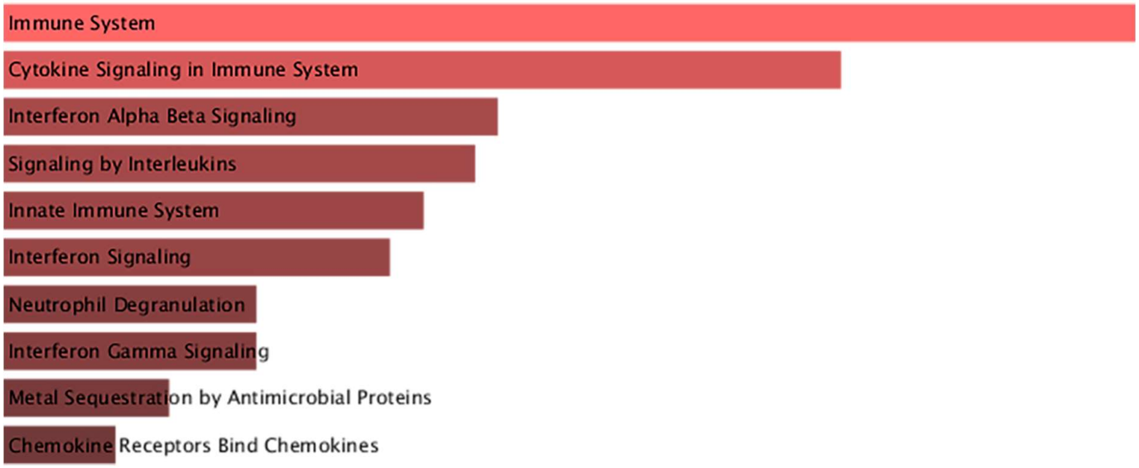
Immune System
Cytokine Signaling in Immune System
Interferon Alpha Beta Signaling
Signaling by Interleukins
Innate Immune System
Interferon Signaling
Neutrophil Degranulation
Interferon Gamma Signaling
Metal Sequestration by Antimicrobial Proteins
Chemokine Receptors Bind Chemokines

b)

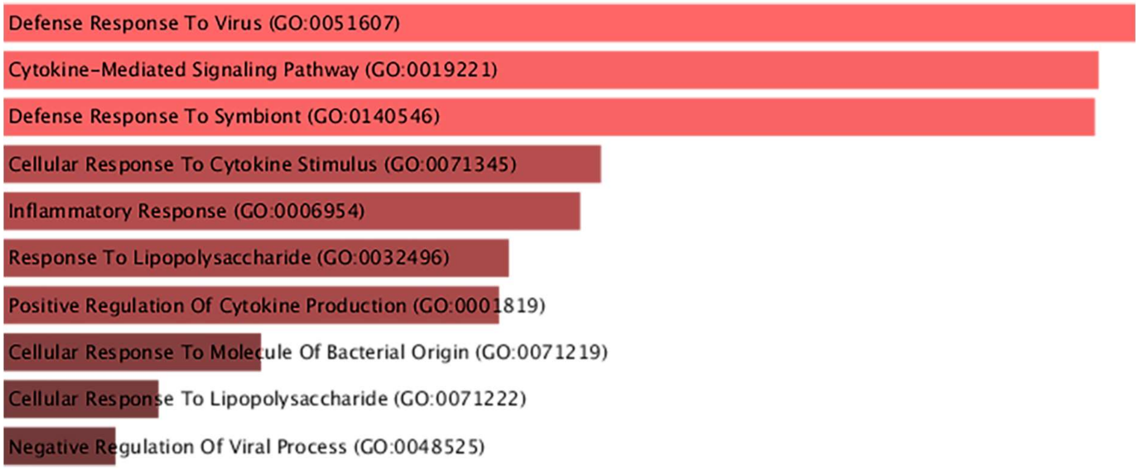
Defense Response To Virus (GO:0051607)
Cytokine-Mediated Signaling Pathway (GO:0019221)
Defense Response To Symbiont (GO:0140546)
Cellular Response To Cytokine Stimulus (GO:0071345)
Inflammatory Response (GO:0006954)
Response To Lipopolysaccharide (GO:0032496)
Positive Regulation Of Cytokine Production (GO:0001819)
Cellular Response To Molecule Of Bacterial Origin (GO:0071219)
Cellular Response To Lipopolysaccharide (GO:0071222)
Negative Regulation Of Viral Process (GO:0048525)

c)

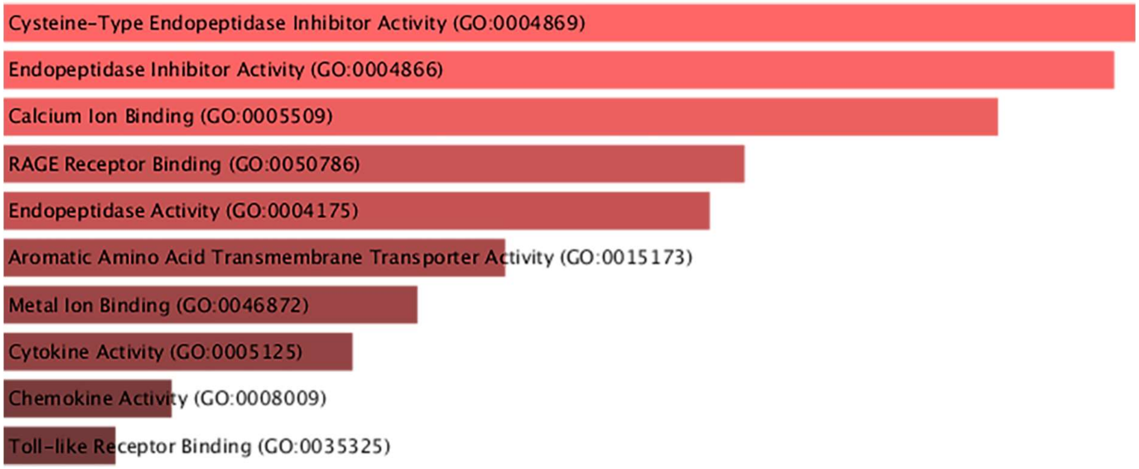
Cysteine-Type Endopeptidase Inhibitor Activity (GO:0004869)
Endopeptidase Inhibitor Activity (GO:0004866)
Calcium Ion Binding (GO:0005509)
RAGE Receptor Binding (GO:0050786)
Endopeptidase Activity (GO:0004175)
Aromatic Amino Acid Transmembrane Transporter Activity (GO:0015173)
Metal Ion Binding (GO:0046872)
Cytokine Activity (GO:0005125)
Chemokine Activity (GO:0008009)
Toll-like Receptor Binding (GO:0035325)

**d)**

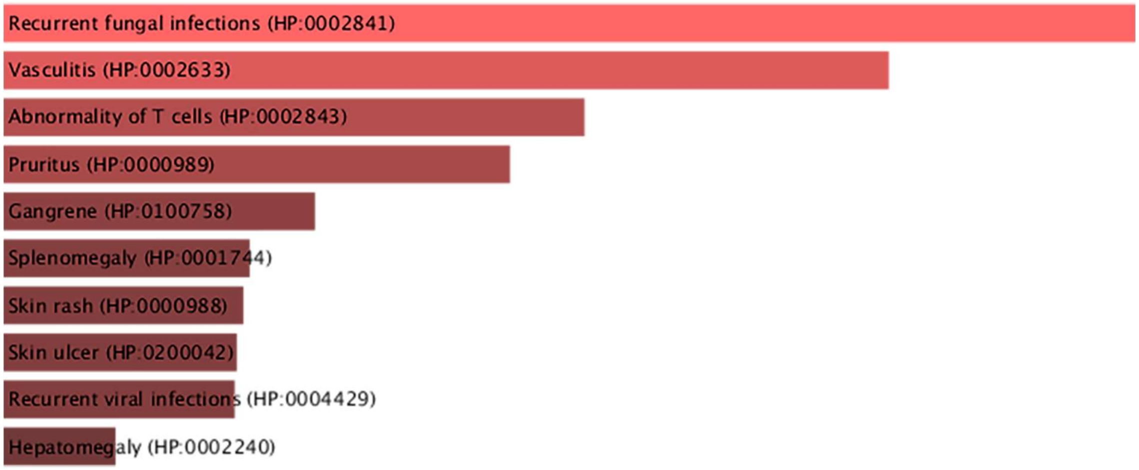


**Supplementary Data Figure 2: Gene Ontology (GO) enrichment analysis of the Meta-Gene Expression Profile (MGEP).** GO enrichment analysis was performed on the 435 DEGs comprising the MGEP using Enrichr. Enriched terms were identified for: **(a)** Pathways, **(b)** Biological Processes, **(c)** Molecular Functions, and **(d)** Human Phenotype Ontology (HPO) Terms. All panels display bar plots of enriched terms, ranked by p-value (descending) to indicate statistical significance.

**a)**

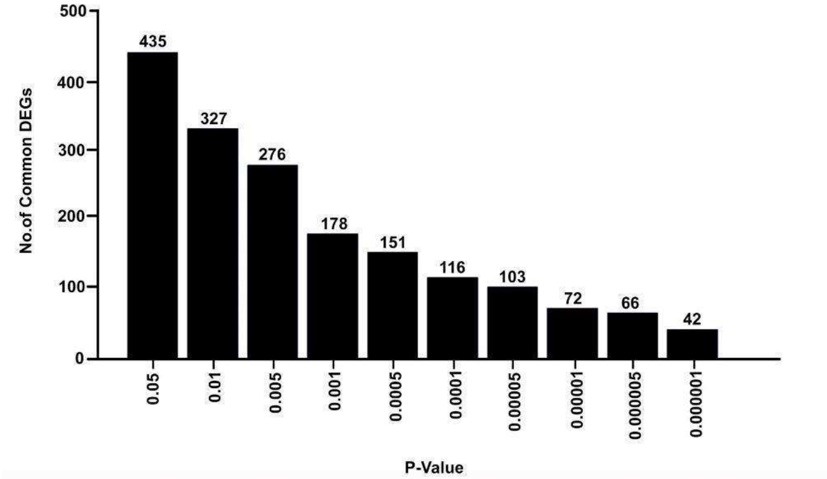

**b)**

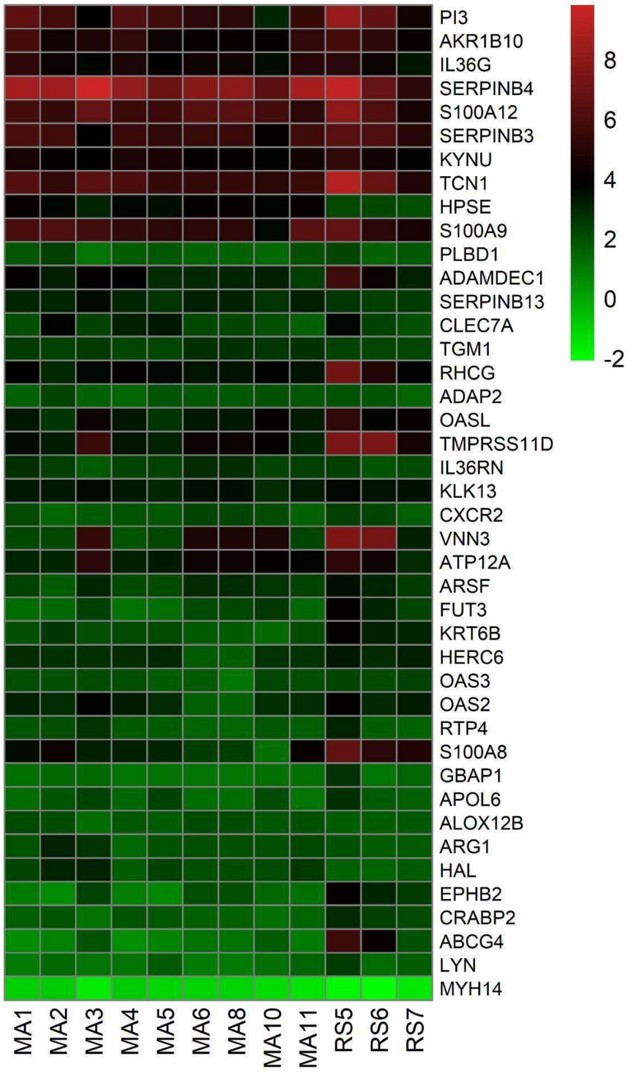


**Supplementary Data Figure 3: Identification and characterization of Highly Significant Meta-Gene Expression Profile (HSMGEP) in psoriasis. (a) Convergence of Differentially Expressed Genes (DEGs) Across Psoriasis Datasets at Increasing Statistical Stringency.** This graph illustrates the cumulative count of shared differentially expressed genes (DEGs) identified across 12 independent psoriasis datasets as a function of increasing statistical significance thresholds. A total of 42 genes met the most stringent significance criterion (q-value < 0.000001), defining the highly significant meta-gene expression profile (HSMGEP), indicating consistent and robust dysregulation across all analyzed cohorts. **(b) Heatmap Depiction of the Highly Significant Meta-Gene Expression Profile (HSMGEP) in Psoriasis.** A heatmap was generated to visualize the expression patterns of the 42 highly significant differentially expressed genes (DEGs) comprising the HSMGEP (q-value < 0.000001). These genes were identified through a meta-analysis of 12 curated psoriasis datasets, encompassing gene expression data from 389 healthy control samples and 312 samples from patients with psoriasis. Notably, *ALOX12B* is included within this 42-gene signature and exhibits a significant upregulation of approximately $\log_2$ fold change, demonstrating its strong and consistent dysregulation in psoriatic skin.

**a)**

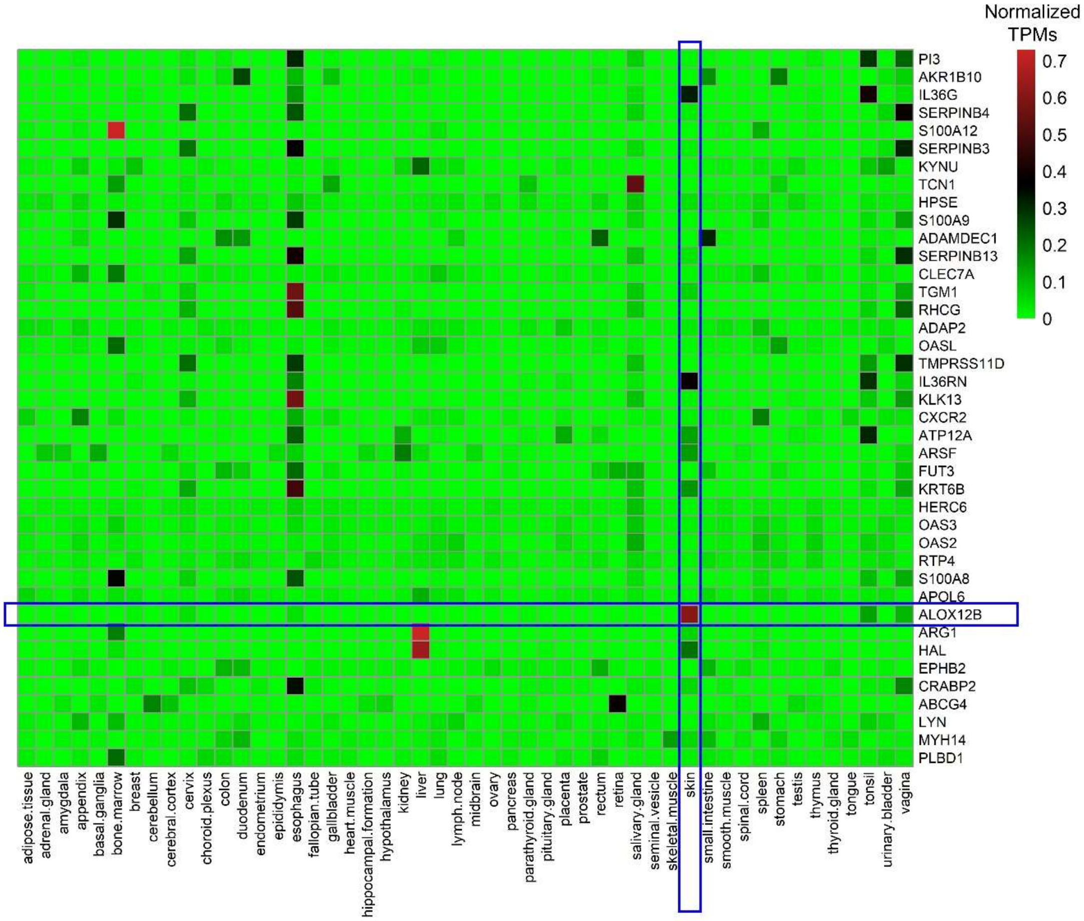


**b)**

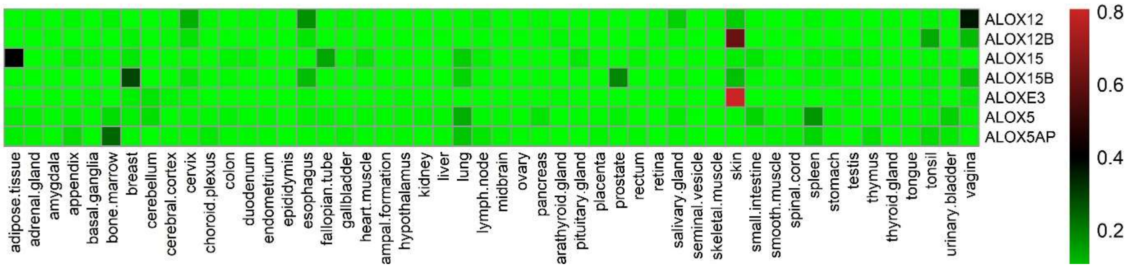


**Supplementary Data Figure 4: Tissue specificity analysis of HSMGEP genes and *ALOX* lipoxygenase family members based on Transcript Per Million (TPM) data from the Human Protein Atlas. (a) Tissue-wide expression profile of Highly Significant Meta-Gene Expression Profile (HSMGEP) genes.** A heatmap was constructed to visualize the normalized TPM expression data obtained from the Human Protein Atlas (HPA). The heatmap demonstrates the relative abundance of transcripts across various human tissues, revealing a pronounced enrichment of *ALOX12B* transcripts in skin tissue, indicating its high tissue specificity. Heatmap shows relative transcript abundance (TPM) across human tissues for 40 of 42 HSMGEP genes (data for 2 genes unavailable in the Human Protein Atlas). **(b) Comparative tissue expression profiles of the arachidonate lipoxygenase (*ALOX*) gene family.** A heatmap was generated to depict the normalized TPM expression profiles of all members of the *ALOX* gene family across a panel of human tissues. The analysis highlights *ALOX12B* and *ALOXE3* as exhibiting significant skin-specific expression, with markedly higher transcript levels in skin compared to other tissues. This finding suggests a specialized role for these two lipoxygenases in skin physiology.

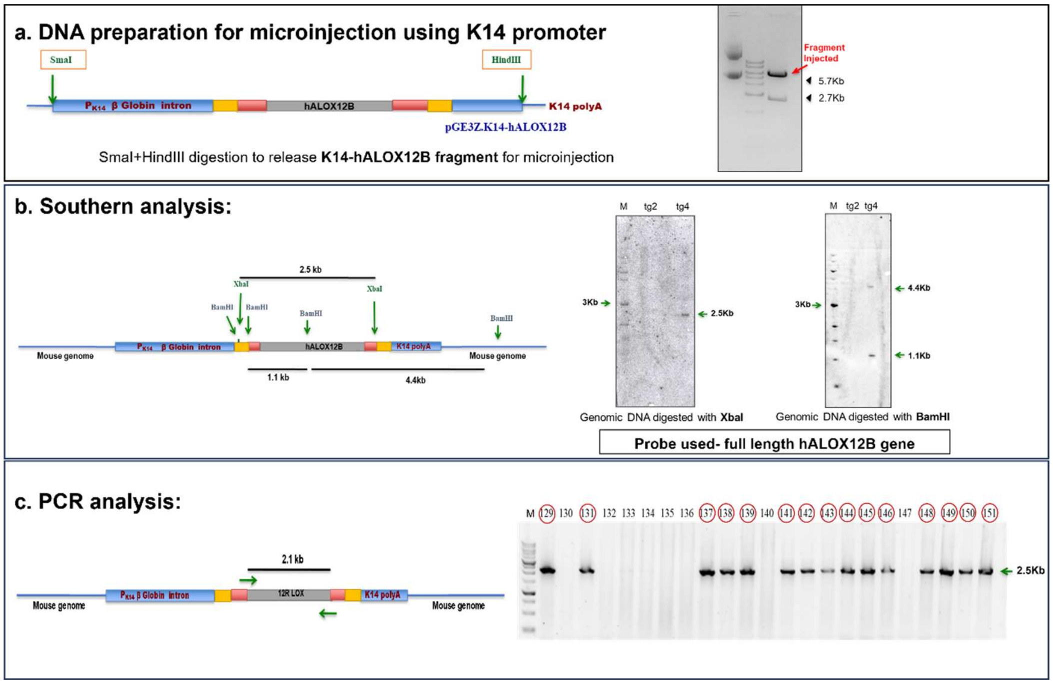


**Supplementary Data Fig. 5: Generation of human ALOX12B overexpressing transgenic mice (Tg-hALOX12B). (a)** Schematic representation of construct preparation for microinjection. To achieve specific overexpression of hALOX12B in the skin, multi-step cloning strategy was implemented. The complete cDNA of the gene was extracted from pCMV6-Neo-hALOX and excised using NotI restriction enzyme digestion. The released cDNA was then used to generate blunt ends for SmaI and blunt-end ligated into the SmaI site of the pBKS plasmid to facilitate the generation of XbaI restriction sites flanking hALOX12B. Further, the gene was released by XbaI from pBKS-hALOX12B and ligated with XbaI in pGE3Z.K14 vector bearing K14 promoter, a beta-globin intron and a polyadenylation (polyA) tail (Data not shown). Finally, the pGE3Z.K14-h12R-LOX vector was digested with SmaI and HindIII to release the entire expression cassette. This cassette includes the K14 promoter, a beta-globin intron, hALOX12B, and a polyadenylation (polyA) tail. The excised fragment was purified and microinjected into mouse embryos at the single-cell stage. **(b)** Genomic DNA was extracted from Tg-hALOX12B, digested with XbaI or BamHI, and subjected to southern blotting. **(c)** Genotyping for the selection of transgenic mice was performed using primers targeting the specific region of hALOX12B by PCR amplification.

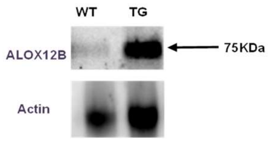


**Supplementary Data Fig. 6**: **ALOX12B is highly expressed in the skin of Tg-hALOX12B.** Representative immunoblots of human ALOX12B in the skin of wild type (WT) and Tg-hALOX12B (TG). Blots represent pooled samples from mice (n = 10).

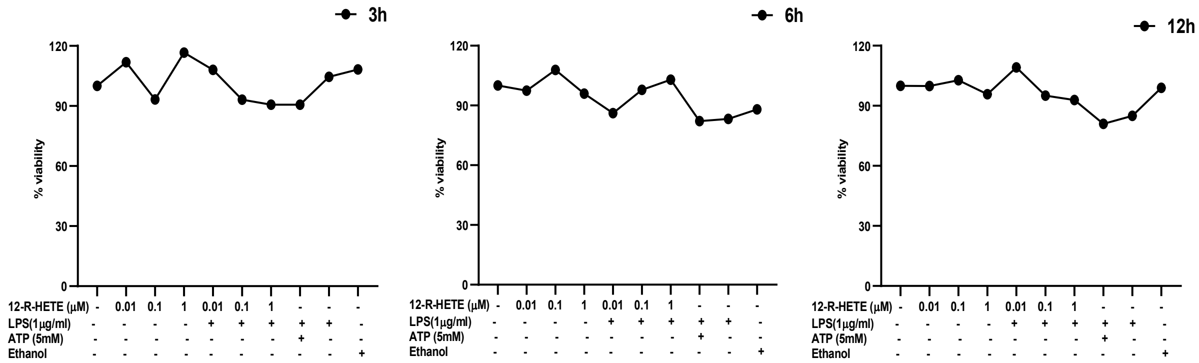


**Supplementary Data Fig. 7: Viability assay represents *12R*-HETE is non-toxic to cells either with or without LPS**. Macrophages treated with different concentrations of 12R-HETE (0.01, 0.1, and 1 µM) with or without LPS (1 µg/ml), ATP (5 mM), and ethanol as vector control are non-toxic when incubated for 3 h, 6 h and 12 h.

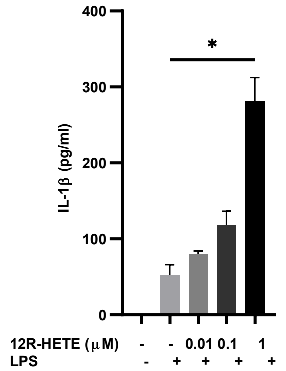


**Supplementary Data Fig. 8: Enhanced levels of active IL-1β in response to *12R*-HETE in macrophages cell line.** Supernatant from the LPS primed (1 µg/ml for 5 h) macrophage cell line treated with different concentrations of 12R-HETE (0.01, 0.1, and 1 µM for 1 h) was assessed for active IL-1β production. The pattern of increase in the levels of active IL-1β production is similar to in bone marrow-derived mice macrophages (BMDM) Fig. 3A.

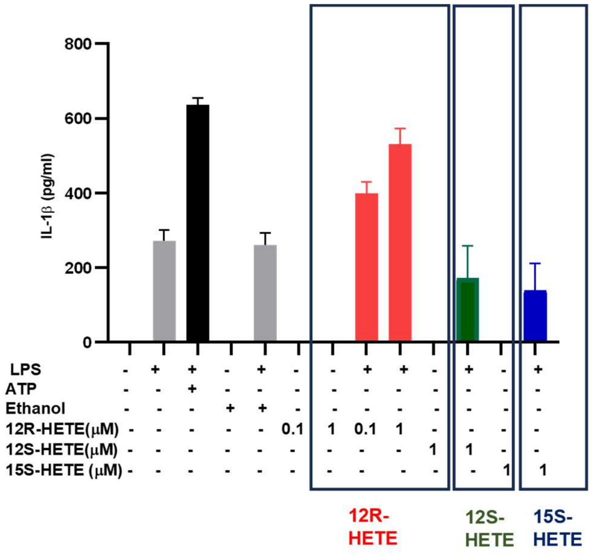


**Supplementary Data Fig. 9: Enhanced level of active IL-1β in LPS primed macrophages is regulated by *12R*-HETE, not the *12S*- and *15S*-HETE.** Active IL-1β production was assessed in the macrophage cell line treated with *12R*-, *12S*-, and *15S*-HETE with or without LPS priming. Results demonstrate the increase in active IL-1β production in macrophages primed with LPS followed by *12R*-HETE treatment as compared to the other controls.

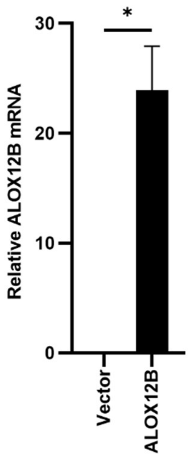


**Supplementary Data Fig. 10: Transcript-level expression of *ALOX12B* in macrophages overexpressing pCMV6-ALOX12B.** Quantitative analysis of *ALOX12B* mRNA expression in macrophages transduced with pCMV6-*ALOX12B* overexpressing constructs.

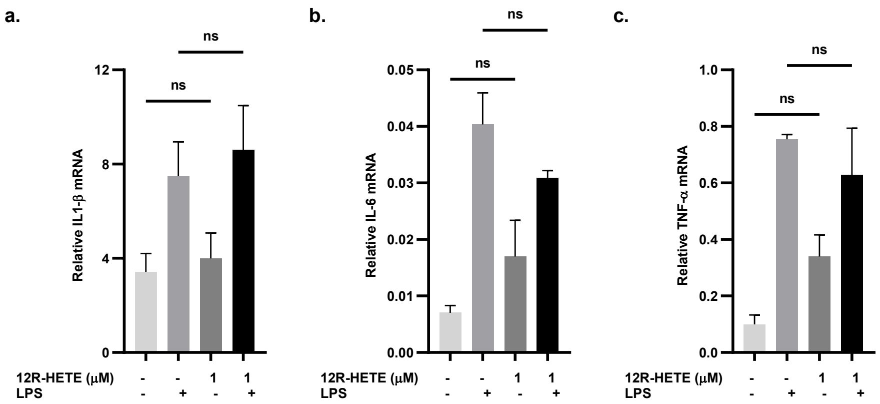


**Supplementary Data Fig. 11:** ***12R*-HETE does not regulate the transcript level expression of IL-1β.** Total RNA was extracted from LPS primed macrophages treated with *12R*-HETE, and performed RT-PCR analysis for the specified genes. The findings did not reveal a notable increase in expression levels of **a.** IL-1β, **b.** IL-6, and **c.** TNF-α when compared with LPS-primed macrophages treated with *12R*-HETE vs LPS primed.

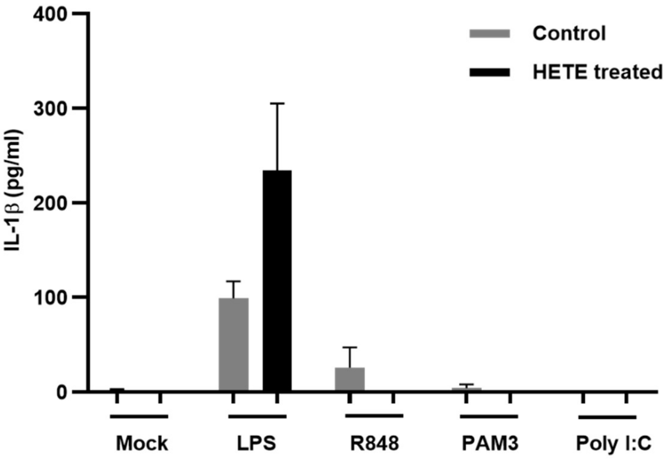


**Supplementary Data Fig. 12:** ***12R*-HETE mediated IL-1β production is predominantly dependent on TLR4 in macrophages.** Macrophages primed with different TLR agonists followed by *12R*-HETE treatment. Active IL-1β was measured through ELISA using cell supernatants.

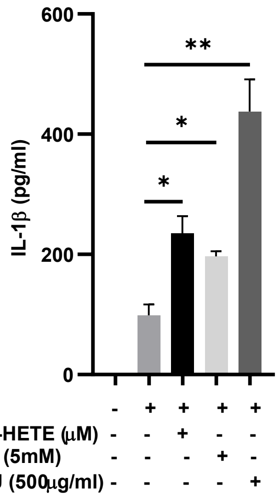


**Supplementary Data Fig. 13: *12R*-HETE acts as a danger signal for the production of active IL-1β from LPS-primed macrophages.** Levels of active IL-1β measured in LPS-primed macrophages treated with different danger signals (ATP and MSU). IL-1β levels in *12R*-HETE treated group is comparable with those treated with ATP or MSU.

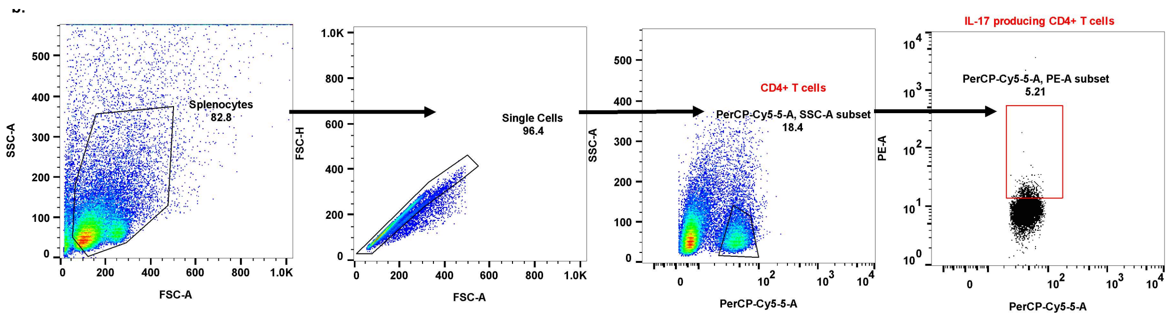


**Supplementary Data Fig. 14: Gating strategy.** The figure depicts the gating strategy employed for the analysis of $CD4^+$ T cells obtained from splenocytes or PBMCs acquired through FACS.

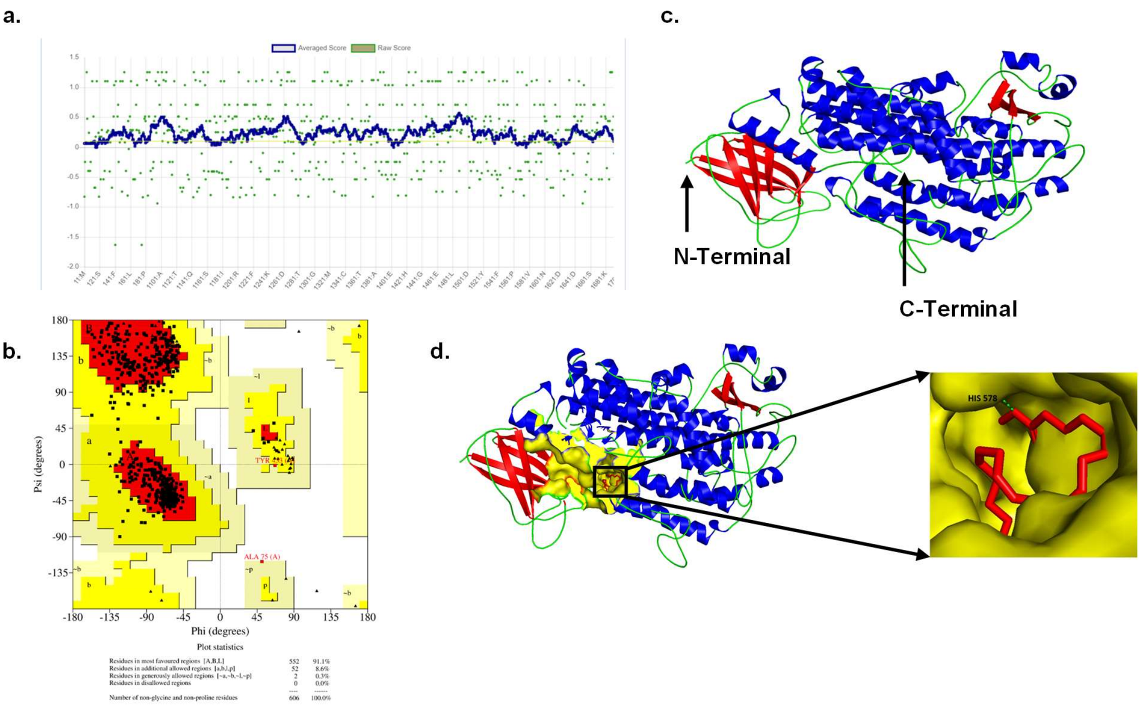


**Supplementary Data Fig. 15: Structural validation of predicted ALOX12B enzyme.** 3D structure of ALOX12B was predicted using tools employed for computational modeling: Phyre2, AlphaFold, and SWISS-MODEL. The amino acid sequence of the enzyme (ID: O75342) was obtained from the UniProt database. Structural validation of ALOX12B through **(a)** Verify 3D **(b)** Ramachandran plot. **(c)** representing the alfa fold structure of ALOX12B indicates C-terminal and N-terminal position. **(d)** ALOX12B structure representing the binding pocket for its substrate (methyl arachidonic acid).

**Supplementary Data Fig. 16: Spectral data of the compounds synthesized through scheme A-D (described in Supplementary methods).**

**(*E*)-3-(5-hydroxy-2-nitrophenyl)-1-(2-hydroxyphenyl)prop-2-en-1-one (3a)**

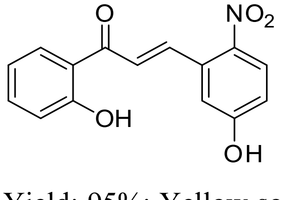


Yield: 95%; Yellow solid; $R_f$ = 0.23 (30% EtOAc/*n*-hexane); $^1$H NMR (400 MHz, $CDCl_3$) δ: 11.90 (s, 1H, ArOH, $D_2O$ exch.), 10.91 (s, 1H, ArOH, $D_2O$ exch.), 8.00 (d, *J* = 9.0 Hz, 1H), 7.91 (d, *J* = 7.9 Hz, 1H), 7.51 (t, *J* = 7.1 Hz, 1H), 7.32 (d, *J* = 2.6, 1H), 6.97-6.91 (m, 2H), 6.82 (dd, *J* = 9.0 & 2.7 Hz, 1H), 5.83 (d, *J* = 7.9 Hz, 1H), 5.68 (s, 1H); MS (ES mass): 286.0 (M+1).

**(*E*)-1-(2-hydroxyphenyl)-3-(2-nitrophenyl)prop-2-en-1-one (3b)**

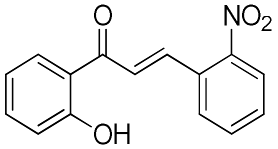


Yield: 65%; Yellow solid; $R_f$ = 0.32 (30% EtOAc/*n*-hexane); $^1$H NMR (400 MHz, $CDCl_3$) δ: 12.56 (s, 1H, ArOH, $D_2O$ exch.), 8.29 (d, *J* = 15.4 Hz, 1H), 8.10 (d, *J* = 8.1 Hz, 1H), 7.90 (dd, *J* = 8.1 & 1.5 Hz, 1H), 7.77-7.69 (m, 2H), 7.60 (t, *J* = 6.9 Hz, 1H), 7.55-7.49 (m, 2H), 7.05 (d, *J* = 8.4 Hz, 1H), 6.95 (t, *J* = 7.1 Hz, 1H); MS (ES mass): 270.1 (M+1).

**2-(5-hydroxy-2-nitrophenyl)-4*H*-chromen-4-one (4a)**

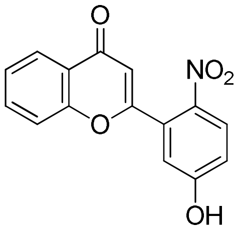


Yield = 95 %; Pale yellow solid; mp: 262-264 oC; Rf = 0.21 (40 % EtOAc/n-hexane); 1H NMR (400 MHz, CDCl3) δ: 11.45 (s, 1H, OH, D2O exch.), 8.16 (d, J = 8.7 Hz, 1H, ArH), 8.08 (dd, J = 7.9 & 1.5 Hz, 1H, ArH), 7.83-7.84 (m, 1H, ArH), 7.57-7.50 (m, 2H, ArH), 7.14-7.09 (m, 2H, ArH), 6.64 (s, 1H, =CH); MS (ES mass): 284.0 (M+1); HPLC: 96.6 %; Column: XBridge C-18 150*4.6 mm, 3.5 µm; Mobile phase: A) 0.1 % TFA in water, B) ACN; (T/B %): 0/10, 3/10, 15/95, 23/95, 25/10, 30/10; Flow rate: 1.0 mL/min; Diluent: ACN: Water (80:20); UV: 230.0 nm; Retention time: 11.3 min.

**2-(2-nitrophenyl)-4*H*-chromen-4-one (4b)**

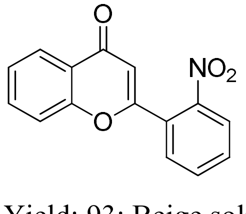


Yield: 93; Beige solid; $R_{f=;}$ Melting Point: ; $^1$H NMR (400 MHz, $CDCl_3$) δ 8.25 (dd, $J_1$ = 2.00 Hz, $J_2$ = 1.60 Hz, 1H), 8.09 (dd, $J_1$ = 0.80 Hz, $J_2$ = 1.60 Hz, 1H), 7.74 (m, 4H), 7.36 (m, 2H), 6.59 (s, 1H); MS (ESI) m/z: 268.10 (M+1); HPLC: 98.78 %, Column: Cosmicsil aura ODS

150*4.6 mm*5µm, Mobile phase A: 5mM $NH_4OAc$ in water, Mobile phase B: $CH_3CN$, gradient (T/%B): 0/10, 20/90, 28/90, 30/10, 35/10; Flow rate: 1.0 ml/min; Diluent: $CH_3CN:H_2O$ (80:20); PDA: 235.0 nm; Retention Time: 14.10 minutes.

**2-(2-hydroxyphenyl)-4*H*-chromen-4-one (4c)**

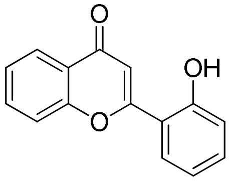


Yield: 92 %; Pale yellow solid; $R_{f=;}$ Melting Point: ; $^1$H NMR (400 MHz, DMSO-$d_6$) δ 10.74 (s, 1H, OH), 8.03 (dd, $J_1$ = 1.20 Hz, $J_2$ = 1.60 Hz, 1H), 7.92 (dd, $J_1$ = 1.60 Hz, $J_2$ = 1.20 Hz, 1H), 7.81 (dd, $J_1$ = 2.00 Hz, $J_2$ = 1.60 Hz, 1H), 7.75 (dd, $J_1$ = 1.20 Hz, $J_2$ = 8.40 Hz, 1H), 7.47 (t, $J$ = 7.40 Hz, 1H), 7.38 (pent, $J$ = 4.3 Hz, 1H), 7.13 (s, 1H), 7.02 (m, 2H); 13C NMR (100 MHz, DMSO-$d_6$) δ 177.7, 161.2, 157.1, 156.3, 134.6, 133.1, 129.1, 125.7, 125.1, 123.6, 119.9, 118.9, 118.1, 117.5, 111.5; MS (ESI) m/z: 239.00 (M+1); HPLC: 98.07 %, Column: Cosmicsil aura ODS 150*4.6 mm*5µm, Mobile phase A: 5mM $NH_4OAc$ in water, Mobile phase B: $CH_3CN$, gradient (T/%B): 0/10, 20/90, 28/90, 30/10, 35/10; Flow rate: 1.0 ml/min; Diluent: $CH_3CN:H_2O$ (80:20); PDA: 210.0 nm; Retention Time: 13.29 minutes.

**2-(5-methoxy-2-nitrophenyl)-4*H*-chromen-4-one (5a)**

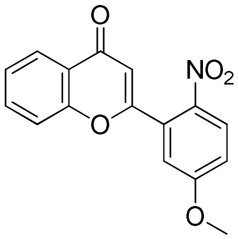


Yield = 92 %; Off-white solid; mp: 183-185 oC; Rf = 0.45 (30 % EtOAc/n-hexane); 1H NMR (400 MHz, CDCl3) δ: 8.25 (d, J = 9.1 Hz, 1H, ArH), 8.10 (dd, J = 7.9 & 1.5, Hz, 1H, ArH), 7.85-7.81 (m, 1H, ArH), 7.58-7.52 (m, 2H, ArH), 7.46 (d, J = 2.8 Hz, 1H, ArH), 7.35 (dd, J = 9.1 & 2.8, Hz, 1H, ArH), 6.75 (s, 1H, =CH), 3.96 (s, 3H, CH3); 13C NMR (100 MHz, CDCl3) δ: 176.8 (C=O), 163.2, 162.4, 155.9, 139.9, 134.7, 129.5, 127.8, 125.9, 124.9, 123.2, 118.3, 117.1, 116.8, 110.9, 56.7 (CH3); MS (ES mass): 298.0 (M+1); HPLC: 99.6 %; Column:

XBridge C-18 150*4.6 mm, 5 µm; Mobile phase: A) 0.1 % TFA in water, B) ACN; (T/B %): 0/5, 20/90, 30/90, 31/5, 35/5; Flow rate: 1.0 mL/min; Diluent: ACN:Water (80:20); UV: 230.0 nm; Retention time: 14.1 min.

**2-(5-ethoxy-2-nitrophenyl)-4*H*-chromen-4-one (5b)**

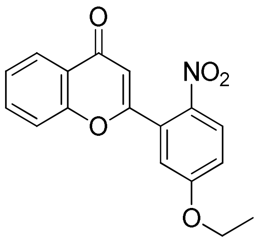


Yield = 90 %; Off-white solid; mp: 153-155 oC; Rf = 0.51 (30 % EtOAc/n-hexane); 1H NMR (400 MHz, CDCl3) δ: 8.23 (d, J = 9.1 Hz, 1H, ArH), 8.10 (dd, J = 7.9 & 1.5, Hz, 1H, ArH), 7.85-7.81 (m, 1H, ArH), 7.58-7.52 (m, 2H, ArH), 7.43 (d, J = 2.8 Hz, 1H, ArH), 7.33 (dd, J = 9.1 & 2.8, Hz, 1H, ArH), 6.74 (s, 1H, =CH), 4.26 (q, J = 6.9 Hz, 2H, CH2), 1.38 (t, J = 6.9 Hz, 3H, CH3); 13C NMR (100 MHz, CDCl3) δ: 176.8 (C=O), 162.5, 163.4, 155.9, 139.8, 134.7, 129.5, 127.8, 125.9, 124.9, 123.2, 118.3, 117.4, 117.1, 110.9, 64.9 (CH2), 14.29 (CH3); MS (ES mass): 312.1 (M+1); HPLC: 99.7 %; Column: XBridge C-18 150*4.6 mm, 5 µm; Mobile phase: A) 0.1 % TFA in water, B) ACN; (T/B %): 0/5, 20/90, 30/90, 31/5, 35/5; Flow rate: 1.0 mL/min; Diluent: ACN:Water (80:10); UV: 230.0 nm; Retention time: 15.2 min.

**2-(2-nitro-5-(prop-2-yn-1-yloxy)phenyl)-4*H*-chromen-4-one (5c)**

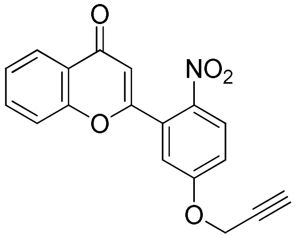


Yield = 90 %; Yellow solid; Rf = 0.48 (30 % EtOAc/n-hexane); 1H NMR (400 MHz, CDCl3) δ: 8.28 (d, J = 9.1 Hz, 1H, ArH), 8.10 (dd, J = 7.9 & 1.5 Hz, 1H, ArH), 7.86-7.81 (m, 1H, ArH), 7.58-7.52 (m, 3H, ArH), 7.42 (dd, J = 9.1 & 2.8 Hz, 1H, ArH), 6.75 (s, 1H, =CH), 5.07 (d, J = 2.4 Hz, 2H, CH2), 3.73 (t, J = 2.4 Hz, 1H, CH); 13C NMR (100 MHz, CDCl3) δ: 176.8 (C=O), 162.1, 160.9, 155.8, 140.6, 134.8, 129.2, 127.7, 125.9, 124.9, 123.2, 118.3, 118.0, 117.5, 110.9, 79.5 (C CH), 77.9 (C CH), 56.7 (CH2); MS (ES mass): 322.0 (M+1).

**2-(2-amino-5-hydroxyphenyl)-4*H*-chromen-4-one (6a)**

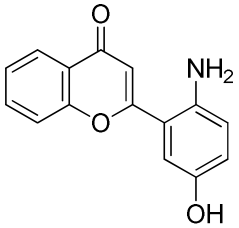


Yield = 85 %; Brick red solid; mp: 202-204 oC; Rf = 0.19 (40 % EtOAc/n-hexane); 1H NMR (400 MHz, CDCl3) δ: 8.82 (s, 1H, OH, D2O exch.), 8.03 (dd, J = 7.9 & 1.4 Hz, 1H, ArH), 7.81-7.77 (m, 1H, ArH), 7.70-7.68 (m, 1H, ArH), 7.49-7.45 (m, 1H, ArH), 6.85 (d, J = 2.5 Hz, 1H, ArH), 6.75-6.69 (m, 2H, ArH), 6.54 (s, 1H, =CH), 5.07 (s, 2H, NH2, D2O exch.); MS (ES mass): 254.1 (M+1); HPLC: 99.8 %; Column: XBridge C-18 150*4.6 mm, 3.5 µm; Mobile phase: A) 0.1 % TFA in water, B) ACN; (T/B %): 0/10, 3/10, 15/95, 23/95, 25/10, 30/10; Flow rate: 1.0 mL/min; Diluent: ACN:Water (80:20); UV: 240.0 nm; Retention time: 7.4 min.

**2-(2-aminophenyl)-4*H*-chromen-4-one (6b)**

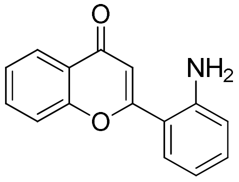


Yield: 88 %; Red solid; $R_{f=}$; Melting Point: ; $^1$H NMR (400 MHz, $CDCl_3$) δ 8.04 (dd, $J_1$ = 1.60 Hz, $J_2$ = 1.20 Hz, 1H), 7.80 (t, $J$ = 4.2 Hz, 1H), 7.72 (dd, $J_1$ = 2.00 Hz, $J_2$ = 8.00 Hz, 1H), 7.47 (t, $J$ = 7.4 Hz, 1H), 7.41 (dd, $J_1$ = 1.60 Hz, $J_2$ = 1.20 Hz, 1H), 7.20 (pent, $J$ = 4.2 Hz, 1H), 6.81 (d, $J$ = 7.6 Hz, 1H), 6.64 (pent, $J$ = 3.90 Hz, 1H), 6.54 (s, 1H), 5.69 (s, 2H, $NH_2$); 13C NMR (100 MHz, DMSO-$d_6$) δ 177.4, 165.4, 156.5, 147.7, 134.4, 132.4, 130.0, 125.7, 125.1, 123.8 119.2, 117.0, 116.5, 115.7, 110.0; MS (ESI) m/z: 238.10 (M+1); HPLC: 89.74 %, Column: Eclips XDB C-18 150*4.6 mm*5µm, Mobile phase A: 5mM $NH_4OAc$ in water, Mobile phase B: $CH_3CN$, gradient (T/%B): 0/10, 20/95, 30/95, 31/10, 35/10; Flow rate: 1.0 ml/min; Diluent: $CH_3CN$:$H_2O$ (80:20); PDA: 240.0 nm; Retention Time: 11.19 minutes.

**2-(2-amino-5-hydroxyphenyl)-6-hydroxy-4*H*-chromen-4-one (6c)**

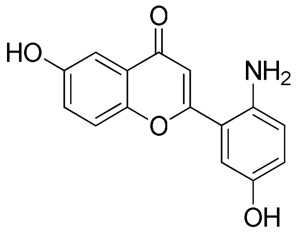

Yield = 80 %; Reddish powder; mp: 130-132 °C; $R_f$ = 0.27 (50 % EtOAc/*n*-hexane); $^1$H NMR (400 MHz, $CDCl_3$) δ: 9.95 (s, 1H, OH, $D_2O$ exch.), 8.79 (s, 1H, OH, $D_2O$ exch.), 7.57 (d, *J* = 8.9 Hz, 1H), 7.32 (d, *J* = 2.9 Hz, 1H), 7.22 (dd, *J* = 8.9 & 3.0 Hz, 1H), 6.84 (d, *J* = 2.4 Hz, 1H), 6.76-6.70 (m, 2H), 6.47 (s, 1H, =CH), 5.04 (broad s, 2H, $NH_2$, $D_2O$ exch.); $^{13}$C NMR (100 MHz, $CDCl_3$) δ: 176.7 (C=O), 164.0, 154.7, 149.6, 148.4, 139.8, 124.2, 122.7, 120.1, 119.8, 118.5, 116.3, 114.2, 108.5, 107.4; MS (ES mass): 270.0 (M+1); HPLC: 92.1 %, Column: X-Bridge Phenyle, 150*4.6 mm, 3.5 µm; Mobile phase: A) 0.05 % TFA in water B) ACN:$H_2O$ (90:10); (T/B %): 0/2, 5/2, 20/90, 25/90, 26/2, 30/2; Flow rate: 1.0 mL/min; Diluent: ACN:$H_2O$ (10:90); UV: 210.0 nm; Retention time: 11.0 min.

**2-(2-amino-5-methoxyphenyl)-4*H*-chromen-4-one (6d)**

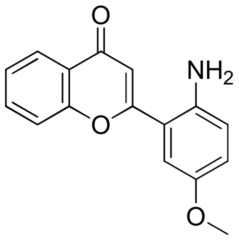

Yield = 90 %; Yellow solid; mp: 118-120 ºC; $R_f$ = 0.38 (40 % EtOAc/*n*-hexane); $^1$H NMR (400 MHz, $CDCl_3$) δ: 8.05 (dd, *J* = 7.8 & 1.4, Hz, 1H, ArH), 7.83-7.78 (m, 1H, ArH), 7.73 (d, *J* = 8.2 Hz, 1H, ArH), 7.49 (t, *J* = 7.1 Hz, 1H, ArH), 7.01 (d, *J* = 2.9 Hz, 1H, ArH), 6.92 (dd, *J* = 8.9 & 2.9, Hz, 1H, ArH), 6.82 (d, *J* = 8.9 Hz, 1H, ArH), 6.64 (s, 1H, =CH), 5.29 (s, 2H, $NH_2$, $D_2O$ exch.), 3.72 (s, 3H, $CH_3$); $^{13}$C NMR (100 MHz, $CDCl_3$) δ: 176.9 (C=O), 164.5, 155.9, 150.6, 141.4, 133.9, 125.3, 124.6, 123.3, 119.8, 118.7, 118.4, 115.5, 112.8, 109.7, 55.5 ($OCH_3$); MS (ES mass): 268.1 (M+1); HPLC: 98.6 %; Column: Eclipse XDB C-18 150*4.6 mm, 5 µm; Mobile phase: A) 0.05 % TFA in water, B) 0.05 % TFA in ACN; (T/B %): 0/10, 5/10, 25/90, 30/90, 31/10, 35/10; Flow rate: 1.0 mL/min; Diluent: ACN:Water (80:20); UV: 240.0 nm; Retention time: 13.4 min.

**2-(2-amino-5-ethoxyphenyl)-4*H*-chromen-4-one (6e)**

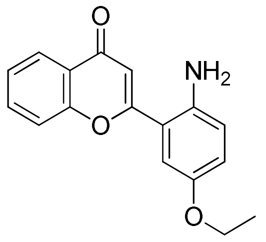

Yield = 90 %; Yellowish brown solid; mp: 88-90 ºC; $R_f$ = 0.4 (40 % EtOAc/*n*-hexane); $^1$H NMR (400 MHz, $CDCl_3$) δ: 8.05 (d, *J* = 6.9 Hz, 1H, ArH), 7.80 (t, *J* = 7.2 Hz, 1H, ArH), 7.73 (d, *J* = 8.3 Hz, 1H, ArH), 7.49 (t, *J* = 7.4 Hz, 1H, ArH), 7.00 (d, *J* = 2.8 Hz, 1H, ArH), 6.91 (dd, *J* = 8.8 & 2.8, Hz, 1H, ArH), 6.80 (d, *J* = 8.9 Hz, 1H, ArH), 6.62 (s, 1H, =CH), 5.28 (s, 2H, $NH_2$, $D_2O$ exch.), 3.97 (q, *J* = 6.9 Hz, 2H, $CH_2$), 1.29 (t, *J* = 6.9 Hz, 3H, $CH_3$); $^{13}$C NMR (100 MHz, $CDCl_3$) δ: 176.9 (C=O), 164.5, 155.9, 149.4, 141.3, 133.9, 125.3, 124.6, 123.3, 120.3, 118.7, 118.4, 115.6, 113.7, 109.7, 63.5 ($CH_2$), 14.75 ($CH_3$); MS (ES mass): 282.1 (M+1); HPLC: 95.4 %; Column: Eclipse XDB C-18 150*4.6 mm, 5 µm; Mobile phase: A) 0.05 % TFA in water, B) 0.05 % TFA in ACN; (T/B %): 0/10, 5/10, 25/90, 30/90, 31/10, 35/10; Flow rate: 1.0 mL/min; Diluent: ACN:Water (80:20); UV: 240.0 nm; Retention time: 14.7 min.

**2-(2-amino-5-(prop-2-yn-1-yloxy)phenyl)-4*H*-chromen-4-one (6f)**

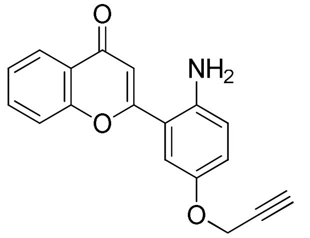

Yield = 90 %; Yellow solid; mp: 153-155 ºC; $R_f$= 0.35 (30 % EtOAc/*n*-hexane); $^1$H NMR (400 MHz, $CDCl_3$) δ: 8.06 ( dd, *J* = 7.9 & 1.6, Hz, 1H, ArH), 7.83-7.79 (m, 1H, ArH), 7.72 (d, *J* = 7.8 Hz, 1H, ArH), 7.52-7.48 (m, 1H, ArH), 7.12 (d, *J* = 2.9 Hz, 1H, ArH), 6.97 (dd, *J* = 8.9 & 2.9 Hz, 1H, ArH), 6.82 (d, *J* = 8.9 Hz, 1H, ArH), 6.62 (s, 1H, =CH), 5.38 (s, 2H, $NH_2$, $D_2O$ Exch.), 4.74 (d, *J* = 2.4 Hz, 2H, $CH_2$), 3.55 (t, *J* = 2.3 Hz, 1H, ≡CH); $^{13}$C NMR (100 MHz, $CDCl_3$) δ: 177.4 (C=O), 164.9, 156.4, 148.8, 142.6, 134.4, 125.8, 125.1, 123.8, 121.4, 119.2, 118.7, 115.8, 115.2, 110.1, 80.1 (C≡CH), 78.5 (C≡CH), 56.7 ($CH_2$); MS (ES mass): 292.1 (M+1); HPLC: 98.8 %; Column: Cosmicsil C18 150*4.6 mm, 5 µm; Mobile phase: A) 0.1 % TFA in water, B) ACN; (T/B %): 0/30, 20/95, 30/95, 31/30, 35/30; Flow rate: 1.0 mL/min; Diluent: ACN:Water (80:20); UV: 240.0 nm; Retention time: 6.6 min.

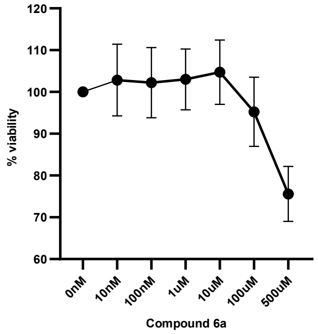


**Supplementary Data Fig. 17: Viability assay of macrophage cell line treated with ALOX12B inhibitor**. Macrophages treated with different concentrations of compound **6a** at a concentration of up to 100 µM, and the percentage of viable cells was determined through MTT assay. Data depicts compound **6a** is non-toxic upto 10 µM when incubated up to 24 h.

**a.** **b.**

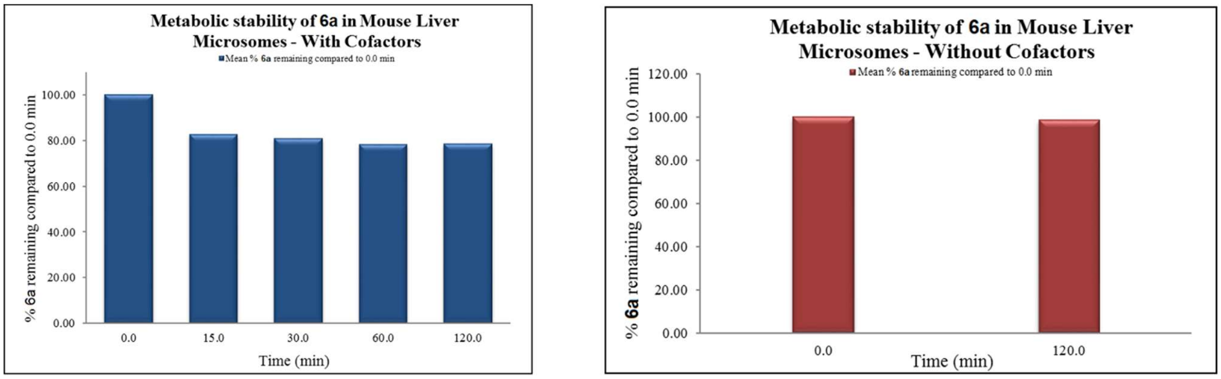


**Supplementary Data Fig. 18: Metabolic stability of 6a in mouse liver microsomes.** Metabolic stability of **6a** in mouse liver microsomes **(a)** with cofactors and **(b)** without cofactors.

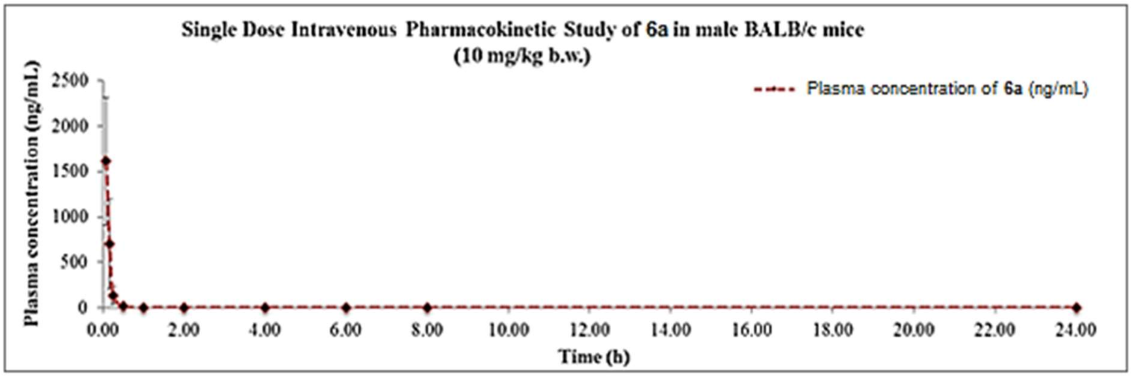


**Supplementary Data Fig. 19: Intravenous pharmacokinetic profile of 6a.** Plasma pharmacokinetics of compound **6a** following a single intravenous dose (10 mg/kg body weight) in mice. Plasma concentrations were quantified using LC-MS/MS, and pharmacokinetic parameters were calculated.

**a.** **b.**

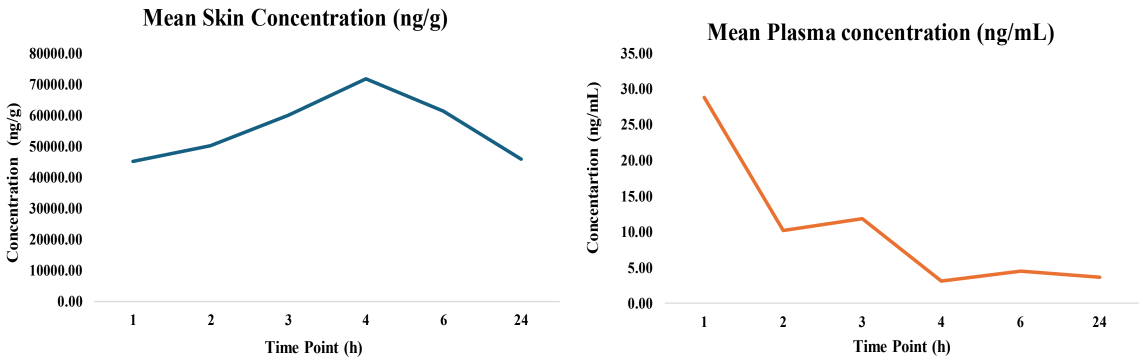


**Supplementary Data Fig. 20: Single dose dermal pharmacokinetics study of 6a in mice showed favourable pharmacokinetic properties.** The plasma and tissue samples were quantified with **6a** till 1.00 to 24.00 h post-dose. **(a)** The mean peak tissue concentration ($C_{max}$) values of **6a** in mice were 71766.33 ng/g at 50 mg/kg/day and **(b)** mean peak plasma concentration ($C_{max}$) values of **6a** in mice were 28.79 ng/mL at 50 mg/kg/day, when measured at regular intervals as mentioned in Table S9.

### Supplementary tables

**Table S1:** Accession numbers (from NCBI GEO) of different datasets representing the total number of healthy and psoriatic skin samples used for metanalysis analysis.

| S.No | DataSet Code | GEO Accession number | No.of healthy skin samples* | No.of Psoriatic samples |
|---|---|---|---|---|
| 1 | D1 | GSE13355 | 64 | 58 |
| 2 | D2 | GSE14905 | 21 | 33 |
| 3 | D3 | GSE78097 | 6 | 13 |
| 4 | D4 | GSE13355[#] | 58 | 58 |
| 5 | D5 | GSE14905[#] | 28 | 33 |
| 6 | D6 | GSE53552 | 24 | 25 |
| 7 | D7 | GSE41662 | 24 | 24 |
| 8 | D8 | GSE30999 | 85 | 85 |
| 9 | D9 | GSE52471 | 13 | 18 |
| 10 | D10 | GSE121212 | 38 | 28 |
| 11 | D11 | GSE67785 | 14 | 14 |
| 12 | D12 | GSE78023 | 14 | 14 |

| | |
|---|---|
| Total no of healthy skin samples* | 389 |
| Total no of unique psoriatic skin samples | 312 |
| Total Samples | 701 |

*(including both the samples from individuals without psoriasis and non-lesional skin from psoriatic patients)
[#](The dataset was duplicated to accommodate two distinct categories of healthy skin samples:
those obtained from individuals without psoriasis and those acquired from non-lesional skin of psoriatic patients)

**Table S2:** The positivity rate of Tg-hALOX12B generation obtained as a result of microinjection experiments where the K14 promoter was used to drive the expression of hALOX12B.

| Experiment | No. of animals used | No. of embryos procured/ Selected for injection. | No. of transfers | Abortions/ Live births | Results |
|---|---|---|---|---|---|
| 1 | 5 | 75/66 | 3 | No/ 7 | Negative |
| 2 | 5 | 124/102 | 5 | 1/23 | Positive (1) |
| 3 | 5 | 135/98 | 4 | 1/23 | Negative |
| 4 | 4 | 73/53 | 2 | Splenomegaly | Negative |
| 5 | 5 | 125/79 | 2 | Died | Negative |
| 6 | 5 | 106/40 | 3 | 1/21 | Negative |
| 7 | 5 | 104/48 | 2 | 31 | Positive (2) |

**Table S3.** List of designed flavone derivatives based on Scaffold-**I** and their respective docking energies with ALOX12B enzyme.

| Flavone derivatives based on Scaffold-I | Docking energy |
|---|---|
| 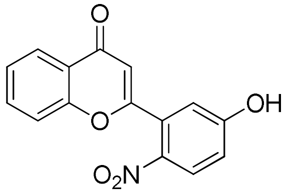 **4a** | -6.67 |
| 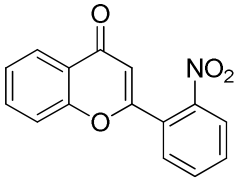 **4b** | -7.01 |
| 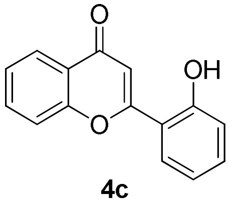 **4c** | -7.0 |

| | |
|---|---|
| 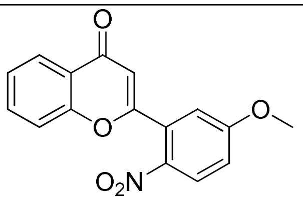 **5a** | -7.19 |
| 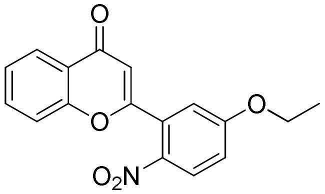 **5b** | -7.29 |
| 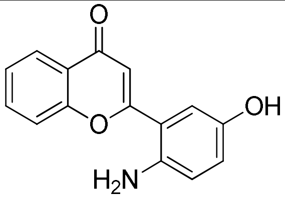 **6a** | -7.19 |
| 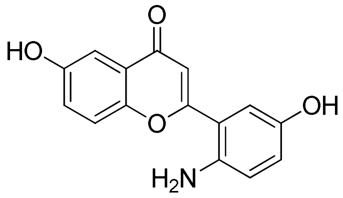 **6c** | -6.96 |
| 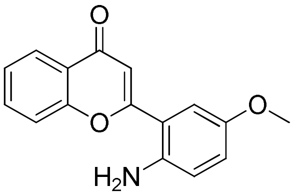 **6d** | -7.31 |
| 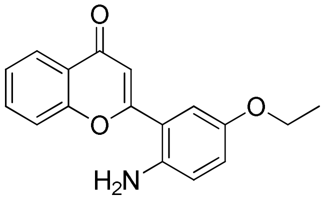 **6e** | -6.93 |
| 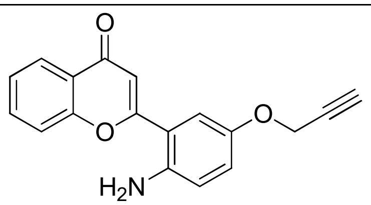 **6f** | -7.41 |

| | |
|---|---|
| 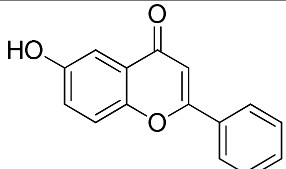 **7** | -6.93 |
| 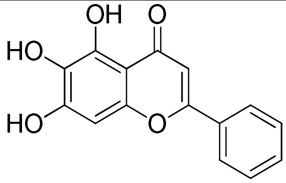 **Baicalein** | -7.04 |

**Table S4.** Percentage inhibition of synthesized flavone derivatives against purified hALOX12B enzyme *in vitro*.

| **Flavone derivatives based on Scaffold-I** | **% hALOX12B inhibition (100 µM)** |
|---|---|
| 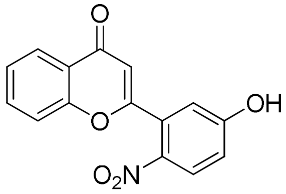 **4a** | 22 ± 6 |
| 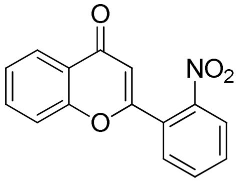 **4b** | No inhibition |
| 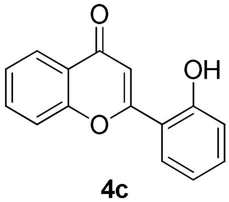 **4c** | No inhibition |
| 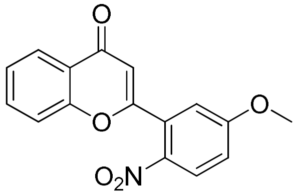 **5a** | No inhibition |

| | |
|---|---|
| 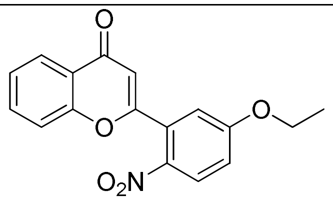 **5b** | No inhibition |
| 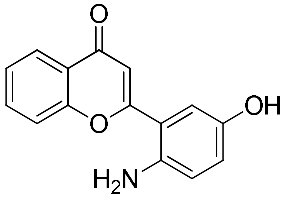 **6a** | 40 ± 10 |
| 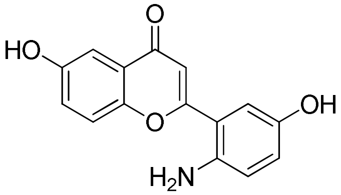 **6c** | 38.33 ± 3.8 |
| 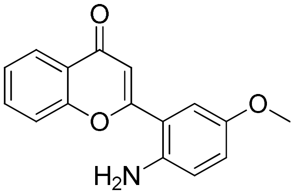 **6d** | No inhibition |
| 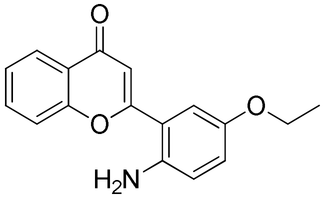 **6e** | No inhibition |
| 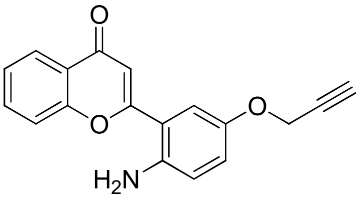 **6f** | No inhibition |
| 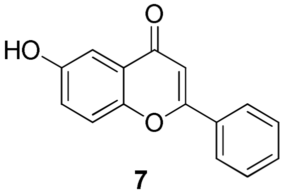 **7** | No inhibition |

**Table S5.** Selective hALOX12B inhibition of compound **6a** over different LOXs.

**a)**

| **% inhibition of ALOX12B enzyme activity** | **Compound 6a** |
|---|---|
| ~ 23 | 10 μM |
| ~ 40 | 100 μM |
| ~ 52 | 200 μM |

**b)**

| **Lipoxygenase enzyme** | **% inhibition of lipoxygenase enzyme activity with Compound 6a (100 μM)** |
|---|---|
| **ALOX12B** | ~ 40 |
| **ALOX12** | ~ 8 |
| **ALOX15** | ~ 24 |
| **ALOX15B** | ~ 6 |

**Table S6.** The half-life ($t_{1/2}$) and *in-vitro* intrinsic clearance of **6a** in mouse liver microsomes were determined through a metabolic stability test in liver microsomes.

| ***In vitro* metabolic stability in mouse liver microsomes** | | |
|---|---|---|
| Test Item | Half-life (min) | Intrinsic Clearance (μl/min/mg of protein) |
| **(6a)** | 495 | 5.6 (Low) |

**Table S7.** Test item administration details of compound **6a** for single dose dermal pharmacokinetic study in mice.

| **Time Point** | **Animal** | **Dose (mg/kg)** | **B.wt** | **Dose (mg)** | **DMSO stock (mg/mL)** | **Dose Volume (mL)** | **Skin Weight (g)** |
|---|---|---|---|---|---|---|---|
| Predose | 1 | 0 | 27.60 | 1.38 | - | 0.138 | 0.75 |
| | 2 | 0 | 31.32 | 1.57 | - | 0.157 | 0.68 |
| | 3 | 0 | 30.52 | 1.53 | - | 0.153 | 0.97 |
| 1 hr | 4 | 50 | 27.41 | 1.37 | 10 | 0.137 | 0.97 |
| | 5 | 50 | 25.86 | 1.29 | 10 | 0.129 | 0.88 |
| | 6 | 50 | 33.77 | 1.69 | 10 | 0.169 | 0.81 |
| 2 hr | 7 | 50 | 27.66 | 1.38 | 10 | 0.138 | 0.68 |
| | 8 | 50 | 28.1 | 1.41 | 10 | 0.141 | 0.63 |
| | 9 | 50 | 30.07 | 1.50 | 10 | 0.150 | 0.88 |
| 3 hr | 10 | 50 | 29.41 | 1.47 | 10 | 0.147 | 0.77 |
| | 11 | 50 | 30.21 | 1.51 | 10 | 0.151 | 0.83 |
| | 12 | 50 | 30.31 | 1.52 | 10 | 0.152 | 0.69 |
| 4 hr | 13 | 50 | 24.12 | 1.21 | 10 | 0.121 | 0.58 |
| | 14 | 50 | 27.24 | 1.36 | 10 | 0.136 | 0.49 |

| | | | | | | | |
|---|---|---|---|---|---|---|---|
| | 15 | 50 | 31.15 | 1.56 | 10 | 0.156 | 0.81 |
| 6 hr | 16 | 50 | 27.4 | 1.37 | 10 | 0.137 | 0.61 |
| | 17 | 50 | 30.51 | 1.53 | 10 | 0.153 | 0.77 |
| | 18 | 50 | 28.99 | 1.45 | 10 | 0.145 | 0.57 |
| 24 hr | 19 | 50 | 30.71 | 1.54 | 10 | 0.154 | 0.61 |
| | 20 | 50 | 26.91 | 1.35 | 10 | 0.135 | 0.57 |
| | 21 | 50 | 28.61 | 1.43 | 10 | 0.143 | 0.61 |

**Table S8.** Clinical signs and mortality observed in single dose dermal pharmacokinetic study of **6a** in Mice.

| Group/ Treatment | Animal Number | Dose | Clinical Sign Observations |
|---|---|---|---|
| Compound (6a) | 01 | 50 mg/kg | Normal |
| | 02 | | Normal |
| | 03 | | Normal |
| | 04 | | Normal |
| | 05 | | Normal |
| | 06 | | Normal |
| | 07 | | Normal |
| | 08 | | Normal |
| | 09 | | Normal |
| | 11 | | Normal |
| | 12 | | Normal |
| | 13 | | Normal |
| | 14 | | Normal |
| | 15 | | Normal |
| | 16 | | Normal |
| | 17 | | Normal |
| | 18 | | Normal |
| | 19 | | Normal |
| | 20 | | Normal |
| | 21 | | Normal |

**Table S9.** Clinical signs and mortality in single dose dermal pharmacokinetic study of **6a** in Mice at different time points.

| Animal ID | Time (Hrs) | Concentration of 6a | | | |
|---|---|---|---|---|---|
| | | Plasma (ng/mL) | Plasma Mean (ng.mL) | Skin (ng/g) | Tissue Mean (ng/g) |
| 1 | Pre-dose | BQL | 0 | BQL | 0 |
| 2 | | BQL | | BQL | |
| 3 | | BQL | | BQL | |
| 4 | 1 | 48.02 | 28.79 | 34663.60 | 45126.25 |
| 5 | | 23.06 | | 35532.84 | |
| 6 | | 15.3 | | 65182.32 | |
| 7 | 2 | 17.38 | 10.19 | 43508.05 | 50212.99 |
| 8 | | 7.09 | | 59229.57 | |
| 9 | | 6.09 | | 47901.35 | |
| 10 | 3 | 3.87 | 11.85 | 51516.43 | 59876.08 |
| 11 | | 7.99 | | 56297.99 | |
| 12 | | 23.68 | | 71813.82 | |
| 13 | 4 | 4.8 | 3.09 | 95139.58 | 71766.33 |
| 14 | | 3.48 | | 73622.24 | |
| 15 | | 0.98 | | 46537.17 | |
| 16 | 6 | 7.13 | 4.46 | 75822.00 | 61257.85 |
| 17 | | 2.65 | | 40238.33 | |
| 18 | | 3.61 | | 67713.22 | |
| 19 | 24 | 5.38 | 3.64 | 54446.25 | 45806.92 |
| 20 | | 4.03 | | 61863.45 | |
| 21 | | 1.51 | | 21111.07 | |

**Table S10.** The $LD_{50}$ and the GSH hazard category of **6a** in Wistar rats to assess the acute dermal toxicity.

| Test Item | $LD_{50}$ (mg/kg body weight) | GSH Classification System | |
|---|---|---|---|
| | | Dermal LD50 (mg/kg body weight) | Hazard Category |
| **6a** | > 2000 | 2000 < ATE ≤ 5000 | 5 |

**Table S11.** GSH classification system*

| Dermal (mg/kg body weight) | Hazard category |
|---|---|
| ATE ≤ 50 | 1 |
| 50 < ATE ≤ 200 | 2 |
| 200 < ATE ≤ 1000 | 3 |
| 1000 < ATE ≤ 2000 | 4 |
| 2000 < ATE ≤ 5000 | 5# |

**The acute toxicity estimate (ATE) for the classification of a test item is derived using the LD50. * Globally Harmonized System of Classification and Labelling of Chemicals (GSH), 9th revised edition, United Nations, New York and Geneva (2021) ST/SG/AC.10/30/Rev.9. # Unclassified**

**Table S12.** Percentage hERG channel (IKr) inhibition of **6a** to assess the cardiac toxicity.

| Concentration (µM) | Max [A] | Mean Current [A] | % Inhibition | Mean % Inhibition | SD |
|---|---|---|---|---|---|
| 30 | 3.10E-10 | 3.10E-10 | 9.05 | 9.05 | 0.60 |
| | 3.08E-10 | | 9.65 | | |
| | 3.12E-10 | | 8.45 | | |
| 10 | 3.01E-10 | 3.04E-10 | 11.84 | 10.88 | 0.86 |
| | 3.05E-10 | | 10.61 | | |
| | 3.07E-10 | | 10.20 | | |
| 3 | 3.02E-10 | 3.05E-10 | 11.53 | 10.55 | 0.86 |
| | 3.07E-10 | | 9.92 | | |
| | 3.07E-10 | | 10.20 | | |
| 1 | 2.82E-10 | 2.93E-10 | 17.34 | 14.09 | 3.27 |
| | 3.05E-10 | | 10.79 | | |
| | 2.93E-10 | | 14.14 | | |
| 0.3 | 3.00E-10 | 2.96E-10 | 12.26 | 13.30 | 1.49 |
| | 2.90E-10 | | 15.01 | | |
| | 2.98E-10 | | 12.62 | | |
| 0.1 | 3.76E-10 | 3.70E-10 | -10.09 | -8.53 | 2.66 |
| | 3.76E-10 | | -10.05 | | |
| | 3.60E-10 | | -5.47 | | |
| 0 | 3.38E-10 | 3.41E-10 | 0.95 | 0.00 | 1.07 |
| | 3.41E-10 | | 0.21 | | |
| | 3.45E-10 | | -1.16 | | |

**Table S13.** Primer sequence.

| S. No. | Target Gene | Primer Sequence |
|---|---|---|
| 1. | hALOX12B for genotyping | FP: GCAGCCCATATGGCCACCTACAAAGTCAGG<br>RP: GCGCTCAAGCTTCTAAATAGAAATGCTGTTCTC |
| 2. | Human ALOX12B transcript | FP: CCATCTCACTGACCATTGTGG<br>RP: CAGGCGGATGATGATGAGC |
| 3. | Human IL-1β | FP: ATGATGGCTTATTACAGTGGCAA<br>RP: GTCGGAGATTCGTAGCTGGA |
| 4. | Human IL-36G | FP: AGGAAGGGCCGTCTATCAATC<br>RP: CACTGTCACTTCGTGGAACTG |
| 5. | Human S100A12 | FP: AGCATCTGGAGGGAATTGTCA<br>RP: GCAATGGCTACCAGGGATATGAA |
| 6. | Human KYNU | FP: GGCTCTCCACCTAGATGAGGA<br>RP: GCTGCTATTTTGGCCCACTTAT |
| 7. | Human HPSE | FP: TCCTGCGTACCTGAGGTTTG<br>RP: CCATTCCAACCGTAACTTCTCCT |
| 8. | Human IL-17A | FP: TCCCACGAAATCCAGGATGC<br>RP: GGATGTTCAGGTTGACCATCAC |
| 9. | Human IL-23A | FP: CTCAGGGACAACAGTCAGTTC<br>RP: ACAGGGCTATCAGGGAGCA |
| 10. | Human IL-22 | FP: GCTTGACAAGTCCAACTTCCA<br>RP: GCTCACTCATACTGACTCCGT |
| 11. | Human IL-18 | FP: TCTTCATTGACCAAGGAAATCGG<br>RP: TCCGGGGTGCATTATCTCTAC |
| 12. | Mice IL-1β | FP: GCAACTGTTCCTGAACTCAACT<br>RP: ATCTTTTGGGGTCCGTCAACT |
| 13. | Mice IL-6 | FP: CTGCAAGAGACTTCCATCCAG<br>RP: AGTGGTATAGACAGGTCTGTTGG |
| 14. | Mice IL-23 | FP: ATGCTGGATTGCAGAGCAGTA<br>RP: ACGGGGCACATTATTTTTAGTCT |
| 15. | Mice IL-17 | FP: TTTAACTCCCTTGGCGCAAAA<br>RP: CTTTCCCTCCGCATTGACAC |
| 16. | Mice TNFα | FP: CCCTCACACTCAGATCATCTTCT<br>RP: GCTACGACGTGGGCTACAG |
| 17. | Mice AIM2 | FP: GTCACCAGTTCCTCAGTTGTG<br>FP: CACCTCCATTGTCCCTGTTTTAT |
| 18. | Mice GAPDH | FP: AAGGTCATCCCAGAGCTGAA<br>RP: CTGCTTCACCACCTTCTTGA |

FP: Forward primer, RP: Reverse primer
